# Divergent impacts of *C9orf72* repeat expansion on neurons and glia in ALS and FTD

**DOI:** 10.1101/2022.11.17.516859

**Authors:** Junhao Li, Manoj K Jaiswal, Jo-Fan Chien, Alexey Kozlenkov, Ping Zhou, Mahammad Gardashli, Luc J. Pregent, Erica Engelberg-Cook, Dennis W. Dickson, Veronique V. Belzil, Eran A Mukamel, Stella Dracheva

## Abstract

Neurodegenerative diseases, such as amyotrophic lateral sclerosis (ALS) and frontotemporal dementia (FTD), are strongly influenced by inherited genetic variation, but environmental and epigenetic factors also play key roles in the course of these diseases. A hexanucleotide repeat expansion in the *C9orf72* (C9) gene is the most common genetic cause of ALS and FTD. To determine the cellular alterations associated with the C9 repeat expansion, we performed single nucleus transcriptomics (snRNA-seq) and epigenomics (snATAC-seq) in postmortem samples of motor and frontal cortices from C9-ALS and C9-FTD donors. We found pervasive alterations of gene expression across multiple cortical cell types in C9-ALS, with the largest number of affected genes in astrocytes and excitatory neurons. Astrocytes increased expression of markers of activation and pathways associated with structural remodeling. Excitatory neurons in upper and deep layers increased expression of genes related to proteostasis, metabolism, and protein expression, and decreased expression of genes related to neuronal function. Epigenetic analyses revealed concordant changes in chromatin accessibility, histone modifications, and gene expression in specific cell types. C9-FTD patients had a distinct pattern of changes, including loss of neurons in frontal cortex and altered expression of thousands of genes in astrocytes and oligodendrocyte-lineage cells. Overall, these findings demonstrate a context-dependent molecular disruption in C9-ALS and C9-FTD, resulting in distinct effects across cell types, brain regions, and disease phenotypes.

**One Sentence Summary:** *C9orf72-associated ALS and FTD* showed a distinct pattern of transcriptome changes, with the largest number of affected genes in C9-ALS in astrocytes and excitatory neurons in upper and deep layers.

## Introduction

Amyotrophic lateral sclerosis (ALS) is a fatal and devastating neurodegenerative disease that manifests in progressive loss of cortical, brainstem and spinal motoneurons resulting in muscle weakness, spasticity, paralysis, and eventually death, typically, within 3-5 years of symptoms onset (*1*). In ∼10% of ALS patients, there is a clear family history of the disease, and pathogenic mutations are found in more than half of these cases (*2*). By contrast, apparently sporadic ALS is considered a complex trait with an estimated heritability of 40–50% (*3*). The pathology and genetics of ALS overlap with another neurodegenerative disease, frontotemporal dementia (FTD), which is the second most common cause of dementia in patients under 65 years (*4*). Aggregates of the RNA-binding transactive response DNA binding 43 protein (TDP-43) accumulate in neurons of almost all ALS cases and approximately half of FTD cases(*5*), and ALS and FTD can occur within the same family and even in the same person (*6*).

Mutations in several genes are causative of ALS, FTD, or ALS/FTD (*7*). The most common genetic cause is a dynamic hexanucleotide (GGGGCC; G_4_C_2_) repeat expansion in the first intron of the *C9orf72* (C9) gene, which is found in ∼40% of familial and ∼12% of all ALS and FTD cases (*8*–*10*). The non-coding G_4_C_2_ expansion appears to have two direct consequences: a loss-of-function that causes *C9orf72* haploinsufficiency, and a gain-of-function associated with the expression of abnormal, bidirectionally transcribed RNAs containing the repeat (*11*). These RNAs accumulate as RNA foci and/or are translated into toxic dipeptide repeat proteins (DPRs) via repeat-associated non-ATG (RAN) translation (*12*–*14*). Notably, ALS- and FTD-associated genes (including *C9orf72)* are expressed in multiple neuronal and non-neuronal cell types in the brain and spinal cord (*15*), raising the question of what impact the mutations have in each individual cell type.

ALS is caused by the progressive degeneration of upper and lower motor neurons, affecting the giant pyramidal cells (Betz cells) in layer 5 of primary motor cortex and the large multipolar alpha motor neurons of the brainstem and spinal cord, whereas FTD results mainly from the degeneration of the large bipolar Von Economo neurons located in in layer 5 of several specific cortical regions (*16*–*18*). Animal and *in vitro* models associated multiple motoneuron-intrinsic (cell-autonomous) pathways with ALS pathogenesis, including glutamate excitotoxicity, mitochondrial dysfunction, axonal transport, RNA metabolism, nucleocytoplasmic transport, and alterations in protein homeostasis (*19*). Although these pathways responded to therapeutic modulation in experimental models, clinical translation has been slow. The only approved therapies for ALS (riluzole, edaravone, and relyvrio) prolong survival by only several months (*20*). This lack of therapeutic progress suggests limitations of current disease models, especially those uniquely focusing on motoneurons. Other neuron types (*21*), as well as astrocytes and microglia, have been suggested to play key roles in the pathogenesis of ALS and FTD (*22, 23*).

We used single-nucleus transcriptomic (snRNA-seq) and epigenomic (snATAC-seq) analysis of human brain samples to investigate the cell-type-specific molecular alterations in ALS or FTD donors with *C9orf72* repeat expansion. In both diseases, we studied two key cortical regions, the motor and frontal cortices, in an attempt to uncover regional differences in disease impact. We detected the most pronounced transcriptional disruption in upper layer excitatory neurons and astrocytes. We also found parallel disruption of the epigenome, with concordant changes in chromatin accessibility, histone modifications, and gene expression in specific cell types. Our comparative analysis of C9-ALS and C9-FTD highlights distinct transcriptional and epigenetic pathologies in these two neurodegenerative diseases.

## Results

### Single nucleus transcriptomes in C9-ALS and C9-FTD specimens identify fine-grained cortical cell types

To investigate the molecular alterations in specific neuronal and non-neuronal cell types from C9-ALS and C9-FTD, we performed snRNA-seq on nuclei isolated from the motor cortex (Brodmann area 4) and dorsolateral prefrontal cortex (Brodmann area 9; hereafter “frontal cortex”) of autopsied C9-ALS (n = 6), C9-FTD (n = 5), and pathologically normal (hereafter “controls”, n = 6) brains (**Table S1**). We obtained 105,120 high-quality nuclear gene expression profiles (45,376 from C9-ALS, 9,445 from C9-FTD, and 50,299 from control subjects), detecting a median of 6,351 unique mRNA molecules and 2,665 genes per nucleus, indicating the high quality of the data. Far fewer nuclei passed quality control in C9-FTD (median 24.3%) compared with controls (66.2%) or C9-ALS (68.4%) (t-test, p < 0.05; **Fig. S1A-C**). This could reflect degeneration in the frontal lobe of FTD patients (*4*), but it could also be caused in part by lower tissue quality of samples from C9-FTD donors.

We used iterative clustering to identify 49 distinct subpopulations of excitatory neurons, inhibitory neurons and non-neuronal cells (**Fig. 1A-B; Table S2**). The clustering was not biased by brain region, donor sex, sequencing batch, individual samples, or diagnosis (**Fig. 1B, Fig. S2**). We annotated these subpopulations based on the expression of known marker genes (**Fig. S3**), which were highly consistent with the cell types annotated in a recent large-scale study of the human motor cortex (**Fig. S4**) (*24*). These results demonstrated the potential resolution of fine-grained cell subtypes our dataset can provide. However, to leverage the information shared across multiple donors when estimating the molecular dysregulation in C9-ALS and C9-FTD, a statistically meaningful number of cells from each donor is required. Therefore, we focused our analysis on 14 major cell types (8 neuronal, 6 glial), defined by grouping subpopulations sharing well-defined markers (**Fig. S3**). At this level of cell type resolution, we observed consistent expression patterns across donors in each major cell type and brain region of the control donors (**Fig. 1C**).

**Fig. 1.**
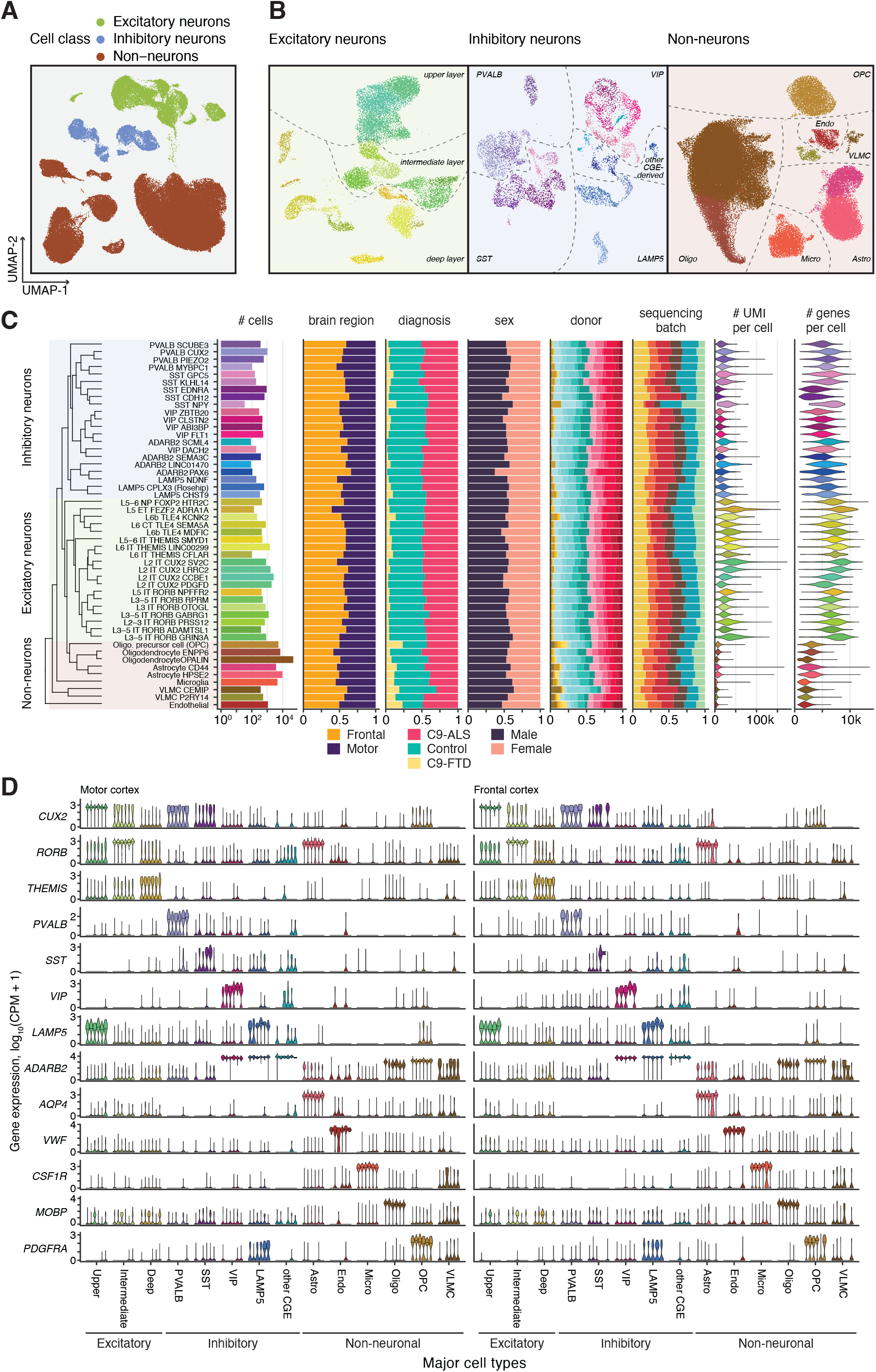
A single cell transcriptomic analysis of human C9-ALS and C9-FTD motor and frontal cortex. (A-B) Identification of transcriptomic cell types from snRNA-seq (n=105,120 cells from 17 donors). Nuclei were first classified into three cell classes in (A), then subpopulations in each class were identified in (B). CGE, caudal ganglionic eminence; Astro, astrocytes; Endo, endothelial cells; Micro, microglia; Oligo, oligodendrocytes; OPC, oligodendrocyte precursor cells; VLMC, vascular leptomeningeal cells; IT, intratelencephalic; CT, corticothalamic; NP, near-projecting; ET, extratelencephalic. See Fig. S3 for marker genes used in the cell type annotation. (C) Neuronal and non-neuronal cell types were distributed across brain regions, donor diagnosis groups, sex, and sequencing batches. UMI, unique molecular identifier. (D) Violin plots show marker gene expression in major cell types in each of the 6 control donors. CPM, counts per million.

### Disruption of gene expression in C9-ALS astrocytes and excitatory neurons

The rare giant pyramidal Betz cells in layer 5b of the human primary motor cortex are considered especially vulnerable in ALS (*25, 26*). Yet, other excitatory and inhibitory neurons, as well as other cortical areas such as the frontal regions, have also been reported to be affected in the disease (*21, 27*). To identify genes dysregulated in C9-ALS in a cell type- and region-specific manner, we compared nuclear transcriptional profiles from C9-ALS and control subjects using a mixed effects model (*28*), while adjusting for demographic and technical covariates (**Methods**).

Excitatory neurons in all cortical layers were profoundly affected in C9-ALS, with hundreds of differentially expressed (DE) genes in each cell type (FDR < 0.05) (**Table S3-4**). Notably, there were more DE genes in upper layer excitatory neurons (L2/3) compared with excitatory neurons in other layers (**Fig. 2A**). In particular, there were 2.48-fold more DE genes in upper-compared with deep-layer (L5/6) excitatory neurons in motor cortex; in frontal cortex, the enrichment was 2.69-fold. This difference was evident even after subsampling each cell type to ensure equal statistical power for detecting DE genes (**Fig. 2B**). We also found that deep layer excitatory neurons were more affected in motor than in frontal cortex, with a larger number of DE genes after controlling for sample size (**Fig. 2B**). In contrast to excitatory neurons, inhibitory neurons had significantly fewer DE genes. Among non-neuronal cells, astrocytes were the most strongly affected in both brain regions (**Fig. 2A-B**).

**Fig. 2.**
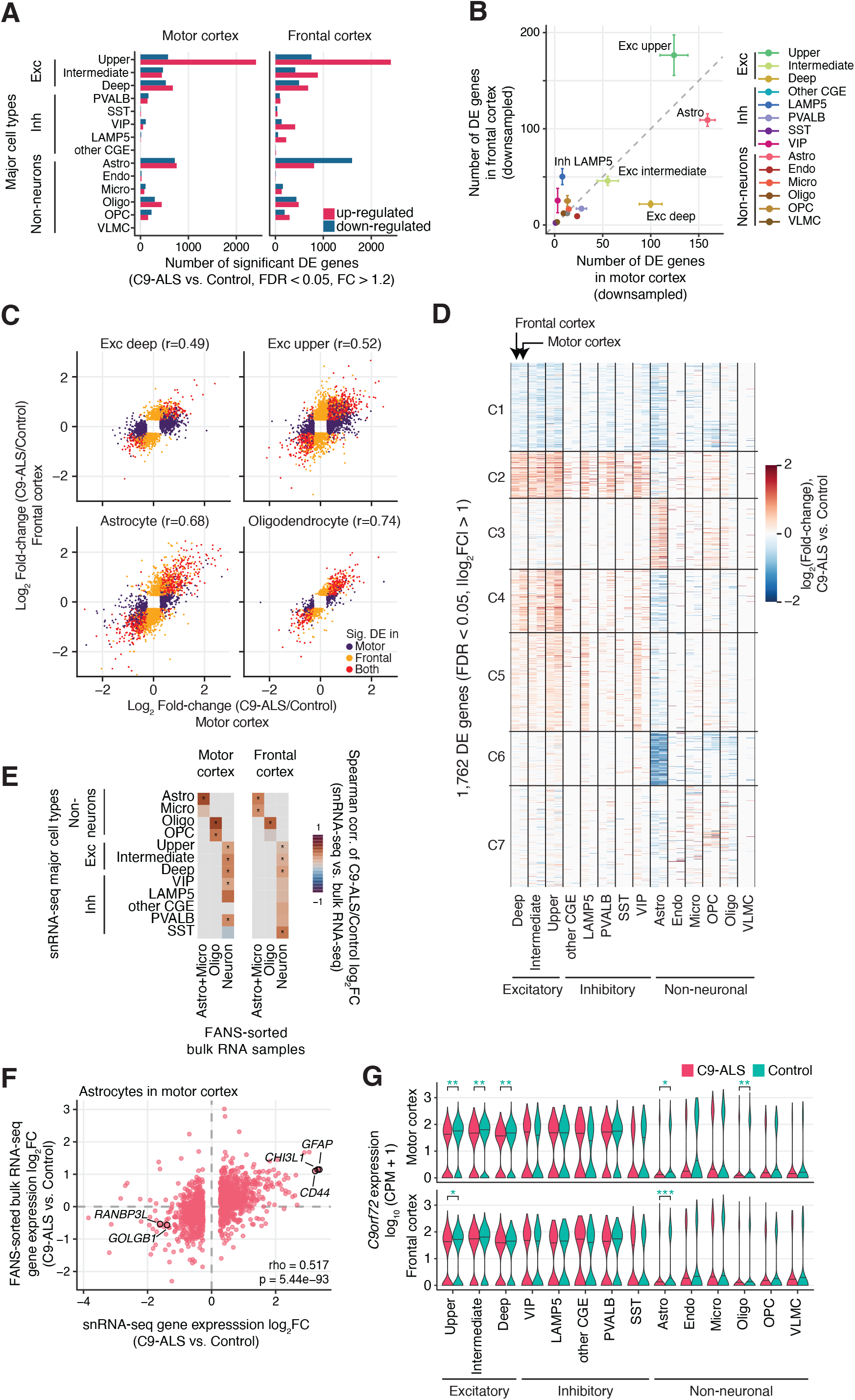
Dysregulation of gene expression in C9-ALS is concentrated in astrocytes and excitatory neurons. (A) The number of differentially expressed (DE) genes (fold-change>1.2, FDR<0.05). Exc, excitatory neurons; Inh, inhibitory neurons. (B) After downsampling to 30 nuclei per donor in each cell type to ensure equivalent statistical power, the majority of DE genes were detected in astrocytes and excitatory neurons. Astrocytes and deep layer excitatory neurons were more affected in motor cortex compared with frontal cortex. Random downsampling was performed 10 times, and the dots and error bars represent mean ± SEM. (C) C9-ALS transcriptional differences were consistent in two cortical regions. r, Pearson correlation coefficient. (D) K-mean clustering analysis of strongly DE genes (>2 fold-change) showed distinct groups of genes are affected in neurons and astrocytes. (E) Validation of snRNA-seq differential expression by bulk RNA-seq in FANS-purified cells. Spearman correlations of the C9-ALS vs. Control FC between the snRNA-seq and bulk-RNA-seq were computed using significant DE genes found in (A). *, Spearman correlation FDR < 0.05. (F) C9-ALS astrocytes exhibit consistent differential expression in snRNA-seq and in bulk RNA-seq. Dots represent significant DE genes (FC > 1.2, FDR < 0.05) found in snRNA-seq. Selected genes of interest with concordant FC are highlighted. rho, Spearman correlation coefficient. (G) *C9orf72* expression was downregulated in C9-ALS in excitatory neurons, astrocytes and oligodendrocytes. Asterisks indicate significant differential expression identified with MAST (*, FDR < 0.05; **, FDR < 0.01; ***, FDR < 0.001). CPM, counts per million.

Overall, transcriptional changes in C9-ALS were highly consistent in motor and frontal cortices (Pearson r = 0.49-0.74, **Fig. 2C**). To characterize the overall pattern of disrupted gene expression across cell types, we performed cluster analysis of the most strongly affected DE genes (fold-change > 2; **Fig. 2D)**. This analysis showed that a unique set of genes was upregulated (cluster C3) or downregulated (C6) in astrocytes compared with other cell types. In contrast, transcriptional changes in neurons were either shared between multiple neuronal subtypes (Clusters C2 and C5) or specific for excitatory neurons (Cluster C4).

We validated our snRNA-seq findings using bulk RNA-seq measurements in purified nuclei from major cortical cell types (**Table S1, Methods**). Using fluorescence-activated nuclear sorting (FANS) with antibodies against pan-neuronal (NeuN) and oligodendrocyte-lineage (SOX10) markers, we purified neurons (NeuN+/SOX10-), oligodendrocyte lineage cells (NeuN-/SOX10+; mature oligodendrocytes and oligodendrocyte precursor cells, OPCs), and non-oligodendrocyte glial cells (SOX10-/NeuN-). The third group mainly consisted of astrocytes and microglia (**Fig. S5A-B; Methods**). Differential gene expression findings were strongly correlated in the single nucleus and FANS RNA-seq data for corresponding cell types in both cortical regions (**Fig. 2E**), especially for astrocytes in motor cortex (Spearman r = 0.517, **Fig. 2F**).

Studies in bulk brain tissues showed that the hexanucleotide repeat expansion in the first intron of *C9orf72* lowers expression of this gene (*9, 29*). Using our single cell data, we found the highest expression of the *C9orf72* gene in neurons, with comparable expression levels in all neuronal subtypes (**Fig. 2G**). *C9orf72* was downregulated in C9-ALS excitatory, but not inhibitory neurons (FDR < 0.05). Among glial cells, only astrocytes and oligodendrocytes showed lower *C9orf72* expression in C9-ALS.

To connect transcriptional alterations in C9-ALS to their impact on protein expression, we used automated Western blotting in bulk motor cortex tissue (**Methods**). We tested multiple antibodies for proteins that were encoded by DE genes. Antibodies against 15 DE gene products consistently detected a protein band of the expected size in control samples. Using these antibodies, we confirmed the predicted changes in protein levels for 6 of the 15 DE genes, with no discordant changes observed for any of the tested proteins (**Figs. 3-4** and **Figs. S6-7**).

**Fig. 3.**
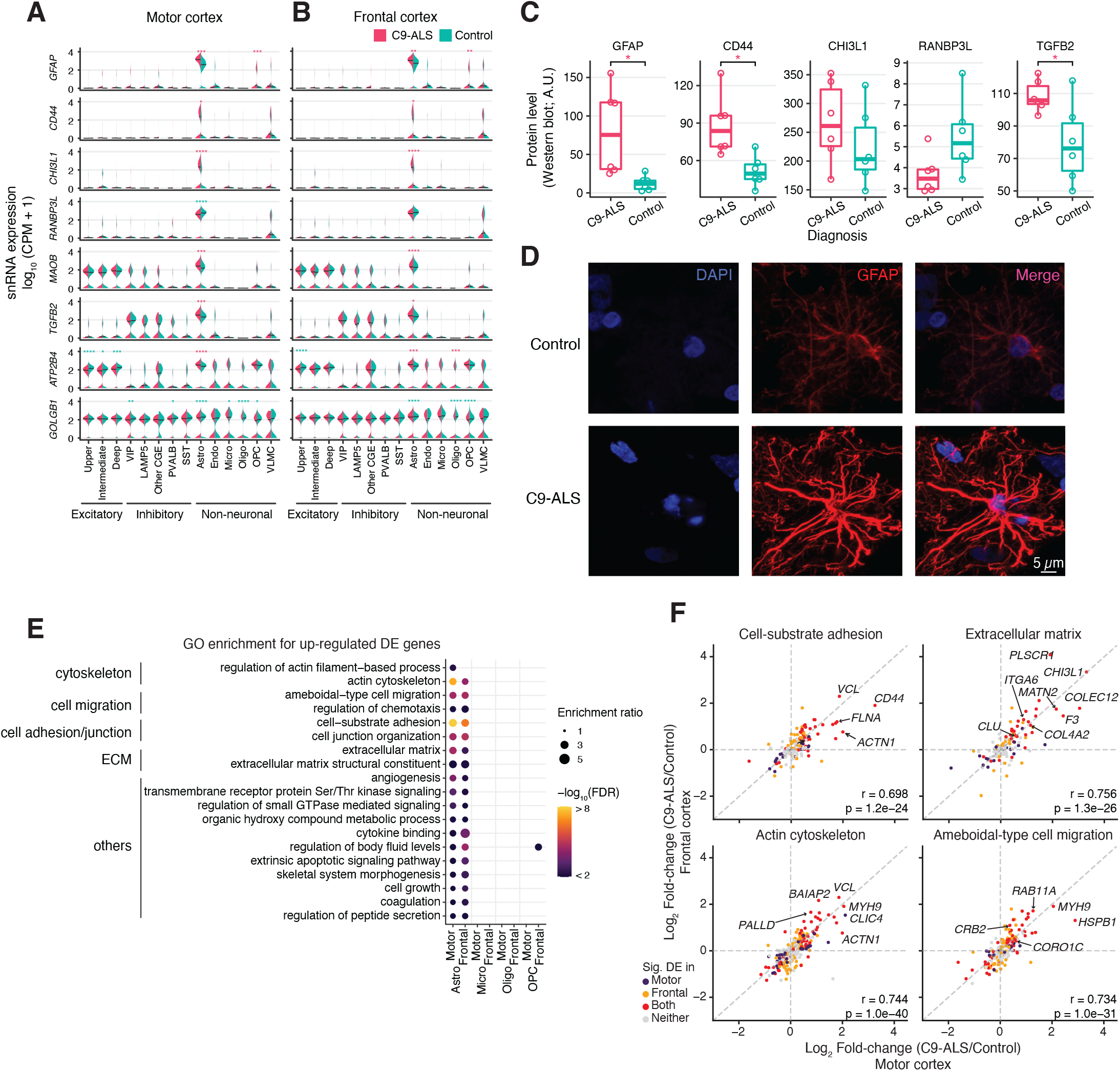
Dysregulation of gene expression in C9-ALS astrocytes suggests activation and structural remodeling. (A-B) Examples of genes that were prominently dysregulated in C9-ALS astrocytes. Asterisks marked the significant upregulated (pink) and downregulated (cyan) DE genes in C9-ALS from the MAST analysis. *, FDR < 0.05; **, FDR < 0.01; ***, FDR < 0.001; ****; FDR < 0.0001. (C) Examples of DE genes in astrocytes with corresponding changes in their protein products. Protein levels were measured by automated Western blot analysis. *, t-test p < 0.05. (D) Representative immunofluorescence images of the GFAP-positive astrocytes in motor cortex (red) obtained using confocal microscopy. DAPI was used to stain nuclei (blue). Immunofluorescence showed strong upregulation of GFAP immunoreactivity in astrocytes in C9-ALS donors. (E) Top Gene Ontology (GO) terms enriched for upregulated DE genes in astrocytes. Enrichment of the same terms for upregulated DE genes in other glia cell types are shown as a comparison. Enrichment ratio is the number of observed genes divided by the number of expected genes from each GO term. ECM, extracellular matrix. See Table S5 for the full list of GO enrichment results. (F) Genes in four functional categories had consistent patterns of differential expression in astrocytes from both brain regions. r, Pearson correlation coefficient.

### Astrocyte transcriptional dysregulation in C9-ALS

We observed a distinct transcriptional signature of astrocyte activation in C9-ALS brains (*30*). mRNA for *GFAP*, a classic marker of astrocyte activation that encodes an intermediate filament protein, increased 10.3-fold in astrocytes of the motor cortex, and 2.4-fold in frontal cortex (**Fig. 3A-B**). Likewise, we found higher expression of the reactive astrocyte-associated genes *CD44* (9.4-fold in motor cortex, 3.7-fold frontal) and *CHI3L1* (10-fold in both regions). CD44-positive astrocytes were found in a variety of neurological diseases (*31*), and elevated levels of chitinases (including CHI3L1) in the cerebrospinal fluid, motor cortex, and spinal cord of ALS patients have been proposed as a biomarker of ALS severity and progression (*32, 33*). We confirmed that GFAP and CD44 proteins were upregulated in C9-ALS bulk motor cortex tissue (p < 0.05; **Fig. 3C; Fig. S6)**. Furthermore, immunofluorescence confirmed that GFAP protein is highly enriched throughout the cell body and processes of astrocytes in C9-ALS motor cortex (**Fig. 3D**).

To assess the functional relevance of the broad pattern of altered gene expression in astrocytes, we performed functional enrichment analysis of DE genes (*34*). In both cortical regions, upregulated genes were enriched for multiple functions including *cell adhesion* and *extracellular matrix* (exemplified by *ITGA6-7, CLU, COL4A2* genes), *actin cytoskeleton* (e.g., *ACTN1, PALLD*), and *cell migration* (e.g., *CRB2, CORO1C*), suggesting substantial cytoskeleton and cell-surface protein remodeling of the C9-ALS astrocytes (GO enrichment FDR < 0.05, **Fig. 3E; Table S5**). DE genes in these functional categories were affected in a consistent manner in both motor and frontal cortices (r > 0.7, **Fig. 3F, Methods**). Remodeling of astrocyte morphology is a hallmark of neurodegeneration (*35*), and our findings highlight the genes involved in this process in C9-ALS. We did not find any functional categories enriched for genes that were downregulated in astrocytes.

We detected changes in the expression of multiple genes whose protein products or functional pathways were previously associated with ALS. For example, several studies reported alterations of transforming growth factor-β2 (TGF-β) signaling in ALS models (*36, 37*). Consistent with these reports, we found strong upregulation of the *TGFB2* gene (1.56-1.95 fold in the two cortices), as well as an increased TGF-β protein level in C9-ALS astrocytes **(Fig. 3A-C, Fig. S6**). Likewise, consistent with the data from *in vitro* and animal ALS models suggesting that endoplasmic reticulum (ER) stress contributes to disease pathogenesis (*38*), *GOLGB1*, which encodes Golgin B1 protein (giantin), was significantly downregulated in astrocytes as well as in oligodendrocyte lineage cells **(Fig. 3A-B**). Giantin is the largest Golgi matrix protein and is involved in the maintenance of Golgi stability and its recovery from ER-to-Golgi transportation-induced stress (*39*). We found that the expression of *RANBP3L*, a gene encoding a Ran-binding protein involved in nucleocytoplasmic transport (*40*), was downregulated 3-fold in motor cortex of C9-ALS astrocytes (**Fig. 3A-B**). We also detected a trend to a reduction of RANBP3L protein in the C9-ALS motor cortex (p = 0.063) (**Fig. 3C, Fig. S6**). This finding is consistent with reports of reduced expression of neuronal nucleocytoplasmic transport proteins in C9-ALS (*41*–*44*).

Our data enabled a granular cell-type-specific analysis of transcriptional alterations in C9-ALS, in some cases, identifying genes with opposite patterns of altered expression in different cell types. For example, we found that *ATP2B4*, which encodes plasma membrane calcium ATPase PMCA4 that catalyzes the hydrolysis of ATP coupled with transport of Ca2+ from the cytoplasm to the extracellular milieu, showed ∼2.0-fold higher expression in C9-ALS *vs*. control astrocytes in both cortical regions (**Fig. 3A-B**). In contrast, *ATP2B4* was downregulated in upper and deep layer excitatory neurons. Astrocytes exhibit dynamic cell state changes mediated by cytosolic Ca2+ (*45*). Thus, increased expression of *ATP2B4* in astrocytes likely represents a protective mechanism to counterbalance upregulation of Ca2+ levels in C9-ALS.

Lastly, recent studies have identified a significant upregulation of monoamine oxidase (MAO) enzyme MAOB in postmortem brains of Alzheimer’s disease (AD) patients (*46*). In addition to the MAO function, MAOB is involved in GAD-independent synthesis of GABA, mediating tonic inhibition in the brain (*47, 48*). *MAOB* was strongly upregulated in C9-ALS astrocytes from motor (3.5-fold) and frontal (2.5-fold) cortices (**Fig. 3A-B**), whereas the homologous enzyme MAOA was downregulated (**Table S3-4**). Thus, our data suggest a role for astrocytic MAOB in modulating the activity of C9-ALS neurons.

### C9-ALS-associated transcriptional alterations in excitatory neurons

Our analysis showed that upper layer (L2/3) and deep layer (L5/6) excitatory neurons were strongly affected in C9-ALS (**Fig. 2B**), and we found a large number of DE genes in both cortical regions (examples in **Fig. 4A; Table S3-4)**. Functional analysis of genes upregulated in these neuronal subtypes in motor cortex showed the most significant enrichments for gene ontology (GO) categories associated with mitochondrial function (*respiratory chain, mitochondrial protein complex, NADH dehydrogenase complex assembly)*, protein synthesis (*protein localization to endoplasmic reticulum, polysome, cytoplasmic translation, translational elongation/initiation*), and cellular proteostasis, the process by which the health of the cell’s proteins is monitored and maintained (*49*) (*protein folding, heat shock protein binding, Golgi-associated vesicle)* (FDR < 0.05) **(Fig. 4C; Table S5)**. There were also notable enrichments for categories associated with nucleocytoplasmic transport (*protein localization to nucleus, RNA localization*), and DNA damage (*nucleoside-excision repair*).

**Fig. 4.**
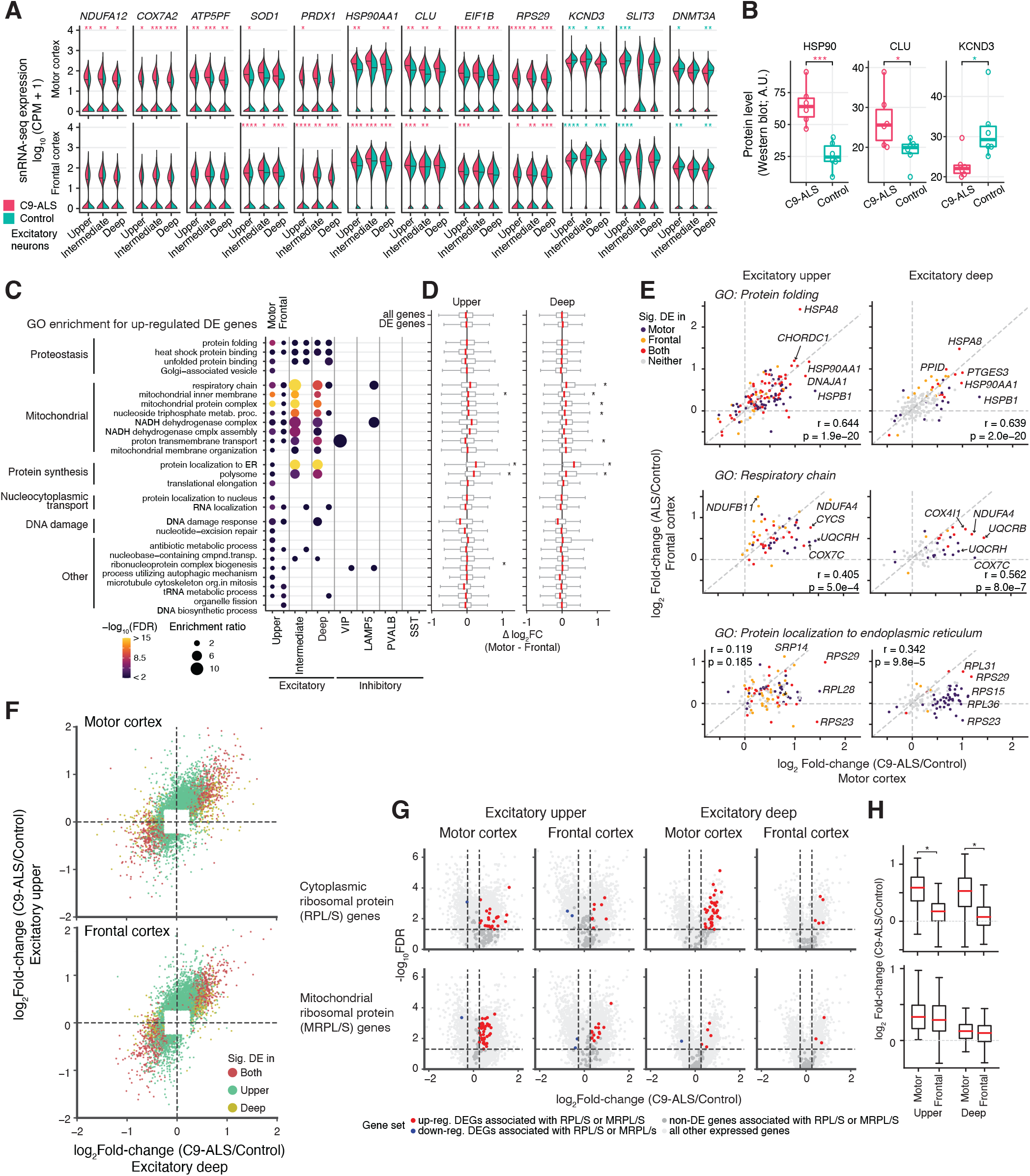
Excitatory neurons have altered expression of metabolic and protein regulatory pathway genes. (A) Examples of genes that were dysregulated in C9-ALS excitatory neurons. Asterisks marked the significant upregulated (pink) and downregulated (cyan) DE genes in C9-ALS from the MAST analysis. *, FDR < 0.05; **, FDR < 0.01; ***, FDR < 0.001; ****; FDR < 0.0001. CPM, counts per million. (B) Examples of DE genes for which corresponding changes in protein abundance were confirmed by automated Western blot analysis. *, t-test p < 0.05; ***, p < 0.001. (C) Top GO terms enriched for genes upregulated in upper and/or deep layer excitatory neurons. Enrichment of the same terms for upregulated DE genes in other neuronal cell types are shown for comparison (also see Table S5). Enriched GO categories (FDR<0.01) were selected by affinity propagation. Enrichment ratio is the number of observed genes divided by the number of expected genes from each GO term. (D) Difference in the fold-change (Δ log_2_FC) between motor and frontal cortex for all expressed genes, all DE genes, and DE genes in each GO category. T-test was used to test whether the Δ log_2_FC in each group of genes were significantly different from the Δ log_2_FC of the “all genes” control set, and asterisks (*) mark the significant differences (FDR<0.001). (E) Comparison of effects in motor and frontal cortices for GO categories exemplifying three major cellular processes enriched for upregulated genes: proteostasis (*protein folding*), mitochondrial function (*reparatory chain*), and protein synthesis (*protein localization to endoplasmic reticulum*). r, Pearson correlation coefficient. (F) Comparison of the C9-ALS vs. control expression fold-changes between DE genes in upper- and deep-layer excitatory neurons. (G) Volcano plots demonstrating the dysregulation of genes associated with cytoplasmic ribosomal proteins (RPL/S, top row) and mitochondrial ribosomal proteins (MRPL/S, bottom row) in upper- and deep-layer excitatory neurons. Dots represent significant upregulated genes (red), downregulated genes (blue), and non-DE genes (dark grey) that are associated with cytoplasmic/mitochondrial ribosomal proteins, and all other expressed genes (light grey). (H) Comparison of the C9-ALS vs. control expression fold-changes of cytoplasmic ribosomal protein genes (top row) and mitochondrial ribosomal protein genes (bottom row) across the neuronal subtypes and cortical regions. *, t-test p-value < 0.05.

Although C9-ALS upper and deep excitatory neurons showed a largely consistent effect on gene expression in motor and frontal cortices (**Fig. 2C**), only a small number of GO categories were enriched for genes upregulated in frontal cortex (**Table S5)**. To understand the regional differences of the disrupted pathways, we compared genes from the enriched GO categories across the two regions (**Fig. 4C-E**; **Methods)**. We found a similar pattern of differential expression for genes associated with cellular proteostasis (*protein folding)* in both regions. However, genes associated with mitochondrial function (*respiratory chain*) were more affected in motor compared to frontal cortex in deep layer excitatory neurons. Genes associated with protein synthesis (*protein localization to endoplasmic reticulum*) were more affected in motor cortex in both neuronal subtypes. This observation was consistent with our downsampling analysis (**Fig. 2B**), showing that certain genes in deep layer excitatory neurons were more strongly affected in C9-ALS in motor cortex compared to frontal cortex. The majority of the DE genes showed stronger upregulation in upper *vs*. deep layer excitatory neurons (**Fig. 4F**).

We detected widespread upregulation of mitochondrial genes in C9-ALS (**Fig. 4C-D)**. Among the DE genes, were those coding for subunits of all four mitochondrial respiratory chain complexes (Complexes I-IV), such as NADH ubiquinone oxidoreductase (*NDUF* family genes), succinate dehydrogenase (*SDHA, SDHB, SDHC*), cytochrome bc1 complex (*UQCR* family and *CYC1*), and cytochrome c oxidase (*COX* family and *CYCS*), as well as ATP synthase complex (*ATP5* family genes) (**Fig. 4A; Table S3-4**). Genes that encode proteins involved in mitochondrial membrane transport (*SLC25A* family, including *SLC25A4*) and chaperones that assist the import of proteins from the cytoplasm into the mitochondrial inner membrane (*TIMM* family, e.g., *TIMMDC1*) were also upregulated. C9-ALS patients had 2.0-2.9 fold higher expression of *SOD1* and *PRDX1*, which encode Cu–Zn superoxide dismutase 1 and peroxiredoxin 1, respectively (**Fig. 4A**). Both enzymes act as scavengers of reactive oxygen species (ROS), which are byproducts of the mitochondrial oxidative phosphorylation (*50*). *SOD1* was the first mutated gene found to be associated with the onset of familial ALS (*51*). Collectively, these findings suggest overall upregulation of mitochondrial function in C9-ALS which leads to increased generation of ROS.

Among upregulated genes associated with proteostasis, there was a large group of genes that encode members of the heat shock protein 70 (HSP70) molecular chaperone family (e.g., *HSPA4, HSPA8, HSPA9*) (**Fig. 4E; Table S3-4**) (*52*). HSP70 proteins are located in major cellular compartments, such as cytoplasm, mitochondria and ER, and are required for aggregation prevention, folding of newly synthesized proteins, conformational maintenance, as well as degradation of misfolded proteins and protein aggregates in autophagic-lysosomal or ubiquitin-proteasome pathways (*53*). HSP70s carry out their functions together with members of large families of co-chaperones. Genes encoding multiple co-chaperones of HSP70 were also upregulated in C9-ALS excitatory neurons, such as members of *DNAJ* and *BAG* families (e.g., *DNAJA1-3, DNAJB1, DNAJC1, BAG2,5,6*) (**Table S3-4**), as well as several other heat shock proteins (e.g., *HSPB1, HSPB11*) that facilitate the function of HSP70s in different protein quality control systems, including the degradation of protein aggregates in a process known as chaperone assisted selective autophagy (*54*–*56*). Furthermore, members of the HSP90 family of chaperons were also upregulated in C9-ALS excitatory neurons (**Fig. 4A, E**). Similar to HSP70, HSP90 chaperones bind to misfolded proteins, including TDP-43, and assist in their transition to native conformation or prevent their accumulation in misfolded state (*57*). Of particular interest is the 1.8-fold increase in expression of *CLU* in upper layer excitatory neurons (**Fig. 4A**). Notably, upregulation of *CLU* was also detected in astrocytes (**Fig. 3F**). *CLU*, which is a risk gene for AD (*58*), encodes a normally secreted chaperone protein, clusterin, that is redirected to the cytosol during ER stress. Clusterin directly interacts with TDP-43 *in vitro* and potently inhibits its aggregation. Thus, increased expression of clusterin could provide an important defense against intracellular proteotoxicity associated with ALS (*59*). We confirmed the upregulation of HSP90 and clusterin proteins in C9-ALS motor cortex (**Fig. 4B; Fig. S7**).

Nearly half of the genes that encode cytoplasmic (*RPL/S* families) and mitochondrial (*MRPL/S* families) ribosomal proteins were upregulated in C9-ALS (**Table S3-4**). In both upper and deep excitatory neurons, we detected a larger number of upregulated cytoplasmic ribosomal genes in motor cortex compared to frontal cortex **(Fig. 4G-H)**. In contrast, this difference was not observed for the mitochondrial ribosomal genes. Other C9-ALS upregulated genes that are associated with protein synthesis were those encoding translation initiation factors (e.g., *EIF1B*) (**Fig. 4A**). These intricate differences could result from region and/or cell specificity of the pathological changes in C9-ALS brain and will require further study.

Genes upregulated in C9-ALS excitatory neurons were linked to processes that are essential for all cell types (e.g., mitochondria function, proteostasis, protein synthesis), whereas genes downregulated in motor cortex were enriched for categories that are specific to neuronal cells (*e*.*g*., *neuronal cell body* category that includes *C9orf72*, and genes that encode several potassium channels) (**Fig. 4A; Fig. S8; Table S3-4**). Downregulated DE genes were also enriched for categories related to neuronal projections (*neuron projection guidance, axon development*), and cell adhesion (*cell-cell adhesion via plasma-membrane adhesion molecules*). Similar to upregulated genes, in both cortical regions, the C9-ALS-associated downregulated genes were more strongly affected in upper *vs*. deep layer excitatory neurons (**Fig. 4F**). Likewise, we did not find significant differences in differential expression between motor and frontal cortices when we compared genes from the GO categories that were enriched in either cortical region (**Fig. S8A-B)**. We also examined the distribution of DE genes across fine-grained excitatory neuron cell types, and found broad enrichment across upper-layer neuron types as well as L6 intra-telencephalic (IT) neurons (**Fig. S8C**).

The downregulated genes include genes that encode brain derived neurotrophic factor (*BDNF*), slit guidance ligands and their receptor (*SLIT1, SLIT3, ROBO2)*, semaphorin proteins and their receptors (*SEMA4B, SEMA4C, SEMA5B, PLXNA1, PLXNA2, PLXNB2*), and RAP1 GTPase activating protein (*RAP1GAP)*. All these proteins are linked to neuronal plasticity in the developing and/or adult brain, and *BDNF* has been implicated in the pathophysiology of several psychiatric and neurodegenerative diseases including ALS (*60*– *65*). Also notable is downregulation of genes that encode several potassium channels (e.g., *KCNN3, KCND3*) (*66*). Unexpectedly, we observed ∼45% decrease in expression of *de novo* DNA methyltransferase *DNMT3A* in the upper- and deep-layer excitatory neurons in motor cortex, suggesting a link to epigenetic dysregulation in C9-ALS. C9-ALS-associated changes in *DNMT3A* levels were previously reported in several studies, yielding conflicting results (*67*–*69*). Consistent with our gene expression data, we detected downregulation of KCND3 protein, as well as a trend for RAP1GAP and DNMT3A proteins in C9-ALS motor cortex (**Fig. 4B; Fig. S7**).

### Disruption of epigenetic regulation in C9-ALS neurons and glial cells

The widespread alterations of gene expression in C9-ALS excitatory neurons, astrocytes, and other cell types raises the question of how such transcriptional changes are established and maintained. We used two complementary assays to determine the epigenetic landscape of C9-ALS brain cells. Single nucleus ATAC-Seq (snATAC-seq) identifies regions of accessible chromatin in single cells (*70, 71*). We generated 109,198 high-quality snATAC-seq profiles (TSS enrichment ≥ 4, unique fragments ≥ 1,000 per cell, **Fig. S9**), and clustered these to generate pseudo-bulk accessibility profiles for 11 major brain cell types (**Fig. S10A**). The snATAC clusters were annotated by transferring labels from the snRNA-seq data (**Table S6; Methods**). The clusters were not driven by brain region, donor sex, individual samples, or diagnosis (**Fig. S10B**), and they had consistent patterns of accessibility at cell type marker genes (**Fig. S10C**). Similar to our snRNA-seq data, we observed fewer nuclei that passed QC in C9-FTD, and C9-FTD nuclei that passed QC also generally had lower TSS enrichment score and number of unique fragments in all major cell types (**Fig. S10D-E**).

We found 39,830 cell-type-specific snATAC peaks marking regions of accessible chromatin (FDR < 0.05, log_2_FC > 0.5 in one cell type vs. the rest; **Fig. 5A**). However, we identified no significant (FDR < 0.05) differential peaks between C9-ALS and control in each major cell type. We also estimated the chromatin accessibility within peaks sharing the same transcription factor (TF) motif with ChromVAR (*72*) **(Fig. S11**). Again, we observed that the variability of ChromVAR scores across nuclei for TF motifs were mainly contributed by the cell type differences. For example, SOX, SPI and TCF were among the topmost variable TF motifs in terms of ChromVAR scores (**Fig. S11A**), and their accessibility were enriched in oligodendrocytes, microglia, and neurons, respectively (**Fig. S11B-D**). Similar to our peak-based analysis, we detected no conclusive changes associated with diagnosis when we directly compared ChromVAR scores between C9-ALS and control nuclei for each TF motif (data not shown).

**Fig. 5.**
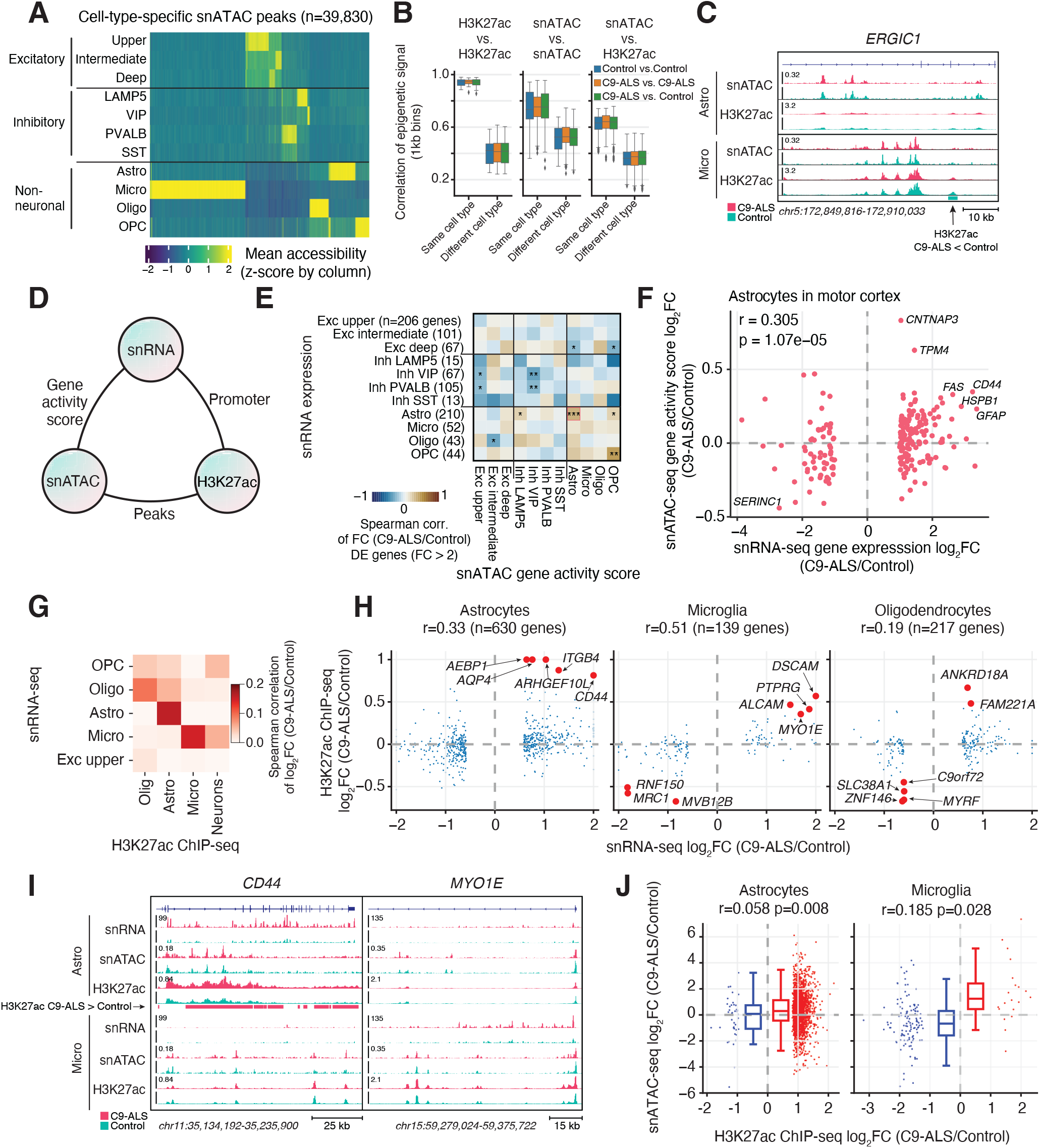
Epigenetic alterations correlate with transcriptome dysregulation in C9-ALS. (A) Normalized chromatin accessibility at cell-type-specific snATAC peaks in 11 major cell types identified in snATAC-seq. (B) Correlation of snATAC and H3K27ac ChIP-seq signal in 1 kb genomic bins. Box-and-whisker plots show the distribution of correlations between replicates for the same cell type, or across different cell types. (C) Genome browser view of snATAC-seq and H3K27ac ChIP-seq signals from astrocytes and microglia at the *ERGIC1* locus. Track height represents pseudo-bulk counts normalized by reads in TSS for snATAC-seq, or average signal (counts normalized to one million reads) across donors for ChIP-seq. Bottom track highlights a H3K27ac peak that is significantly reduced in C9-ALS. (D) Schematic of pairwise comparisons (shown in subsequent panels) of C9-ALS effects on the transcriptome (snRNA-seq), chromatin accessibility (snATAC-seq), and histone modification H3K27ac (ChIP-Seq). For each pair of data modalities, we correlated the fold-change of gene- or H3K27ac peak-associated signals. (E-F) Comparison of snRNA with snATAC. (E) Spearman correlation of the C9-ALS vs. control fold-change (FC) for snRNA expression vs. snATAC gene activity score in motor cortex. The analysis was limited to strongly DE genes (FC > 2) in each major cell type in motor cortex. *, p < 0.05; **, p < 0.01; ***, p < 0.001. See Fig. S10F for frontal cortex data. (F) Scatter plot illustrating the significant correlation between differential gene expression (snRNA-seq) and snATAC-seq changes in astrocytes in motor cortex. Selected genes with high concordant FC are labeled. r, Spearman correlation coefficient. (G,H) Comparison of snRNA with H3K27ac ChIP-Seq, showing Spearman correlation (G) and scatter plots (H) of DE gene expression (snRNA-seq) vs. promoter H3K27ac signal (ChIP-seq). Genes with concordant and biggest H3K27ac signal fold-changes were highlighted in red. (I) Browser view of the *CD44* and *MYO1E* loci, showing the correspondence of epigenomic and transcriptomic signals. Track height represents average RPKM across donors for snRNA-seq, pseudo-bulk counts normalized by reads in TSS for snATAC-seq, or average signal (counts normalized to one million reads) across donors for ChIP-seq. Pink rectangles highlight significant H3K27ac differential peaks that were increased in astrocytes from C9-ALS samples. (J) Correlation of snATAC vs. H3K27ac ChIP-seq signal at differential H3K27ac peaks. Box plots show the distribution for upregulated and downregulated peaks.

Although snATAC-seq can in principle provide fine-grained resolution of cell types, the accuracy of clustering and thus the reliability of the pseudo-bulk profiles is limited by the sparse data from individual cells. Indeed, we found that the distinction between neuronal cell types was not as clear using snATAC-seq data (**Fig. S10A, C**) as it was using snRNA-seq (**Fig. 1A**). We therefore used a complementary strategy, isolating bulk samples of nuclei from four major populations by FANS. Using antibodies against three cell-type-specific nuclear markers, we purified astrocytes (NeuN–/SOX10–/IRF5–), oligodendrocytes (NeuN–/SOX10+/IRF5–), microglia (NeuN– /SOX10–/IRF5+), and neurons (NeuN+/SOX10–/IRF5–) (**Fig. S5B**). We then profiled active enhancers and promoters marked by histone 3 lysine 27 acetylation (H3K27ac) using ChIP-seq (**Fig. S5C**) (*73*). The ChIP-seq data had very high signal/noise ratio, with strong correlation between replicates for the same cell type (r=0.94±0.029, mean ± s.d.) and lower correlation for different cell types (r=0.41±0.084; **Fig. 5B**). The correlation was equally strong between ALS and control subjects as it was between control subjects. snATAC data had lower inter-replicate correlation for the same cell type (r=0.76±0.16) and higher correlation between cell types (r=0.52±0.11; **Fig 5B**).

We observed that the ChIP-seq data correlated strongly with the snATAC-seq tracks from corresponding cell types (r=0.62±0.086), while the correlation for different cell types was lower (r=0.36±0.087) (**Fig. 5B**). Both assays marked cell-type-specific regions of accessible and active regulatory chromatin. The correspondence between H3K27ac and snATAC data at cell-type-specific enhancers was evident at the locus of the ALS risk gene *ERGIC1 (74)*, where multiple snATAC peaks coincide with H3K27ac peaks (**Fig. 5C**). The ChIP-seq tracks generated from bulk nuclei had deeper coverage than the snATAC-seq data (on average 2.25-fold more unique mapped sequence fragments), and we observed a corresponding higher signal/noise ratio in the H3K27ac tracks (**Fig. 5C**).

We found multiple differentially acetylated (DA) chromatin regions in C9-ALS subjects compared with controls (FDR < 0.05; **Table S7**). There was a large difference in numbers of DA regions between the cell types, with 16 DA regions detected in neurons vs. 2945 in astrocytes. For example, we observed more active chromatin (increased H3K27ac), along with increased gene expression, at the *CD44* locus in astrocytes from C9-ALS compared to control subjects (**Fig. 5I**). This disparity is likely explained by heterogeneity of neuronal subtypes which precludes the detection of the DA regions in the bulk population. Notably, as stated above,differential accessibility analysis of the snATAC-seq datasets did not reveal significant differences between ALS and control subjects even in astrocytes. We attributed the lack of differential snATAC-seq peaks to the lower signal/noise ratio in these datasets, as well as the statistical burden of multiple comparisons in analysis of the thousands of genome-wide open chromatin regions.

To further characterize patterns of disease-related differences in epigenetic regulation and connect them with transcriptional alterations, we investigated genome-wide correlations among all three data types: H3K27ac ChIP-Seq, snATAC-seq, and snRNA-seq. In particular, we examined the relationship between cell-type-specific signatures of ALS in each pair of datasets using gene expression, gene activity score, and H3K27ac peaks (**Fig. 5D**). To link gene expression with open chromatin (snRNA vs. snATAC), we calculated the gene activity score for strongly DE genes (FC > 2) (**Methods**). Disease-associated changes in gene expression correlated with open chromatin for astrocytes and OPCs in motor cortex, and in astrocytes and microglia in frontal cortex (FDR < 0.05, **Fig. 5E, Fig. S10F**). Indeed, for astrocytes we found that key upregulated markers including *GFAP, CD44, HSPB1, TPM4*, and *CNTNAP3*, contained more open chromatin in C9-ALS subjects (**Fig. 5F**). By contrast, significantly downregulated genes such as *SERINC1* contained less open chromatin. Notably, we did not observe a significant correlation between snATAC and snRNA signatures of C9-ALS neurons, oligodendrocytes, or microglia, potentially indicating that chromatin accessibility is less affected in those cell types.

Next, we compared differential expression with chromatin activation at the gene promoter (snRNA vs. H3K27ac). This analysis revealed a strong positive correlation for three glial cell types (astrocytes, oligodendrocytes, and microglia; p < 10-5, **Fig. 5G-H**). The correlation was especially strong for microglia, even though fewer genes were DE in microglia compared with astrocytes (r=0.51 for microglia and r=0.33 for astrocytes). The strong consistency of C9-ALS-related differential snRNA and H3K27ac signals in microglia supports the robustness of our observations in these cells. For example, we found that the gene *MYO1E* was transcriptionally upregulated in microglia and had a corresponding enrichment of active chromatin at its promoter (**Fig. 5H-I**). Whereas ALS-related differences in expression were correlated with H3K27ac in a cell-type-specific manner for the non-neuronal cells (**Fig. 5G**), the neuronal ChIP-seq data was less consistent with snRNA, most likely because it represents a mixture of diverse neuronal subtypes.

Finally, we directly compared the two epigenetic data types for the glial cells (**Fig. 5J**), focusing on H3K27ac differentially acetylated peaks. We found that C9-ALS-related changes in chromatin accessibility were positively correlated with H3K27ac in both astrocytes and microglia (p < 0.05, **Fig. 5J**).

### A distinct signature of glial dysregulation in C9-FTD

Our initial analyses showed striking differences in snRNA-seq data from C9-FTD compared with C9-ALS, despite the shared genetic risk factor. Notably, excitatory and inhibitory neurons were depleted 2.1-fold specifically in the frontal cortex in C9-FTD, but not C9-ALS (**Fig. 6A-B**). Neurons were also depleted in many of the motor cortical samples, but the proportion was more variable and not significantly different from controls on average (**Fig. 6B**). Among neurons, the ratio of excitatory to inhibitory neurons was not significantly different (**Fig. 6C**).

**Fig. 6.**
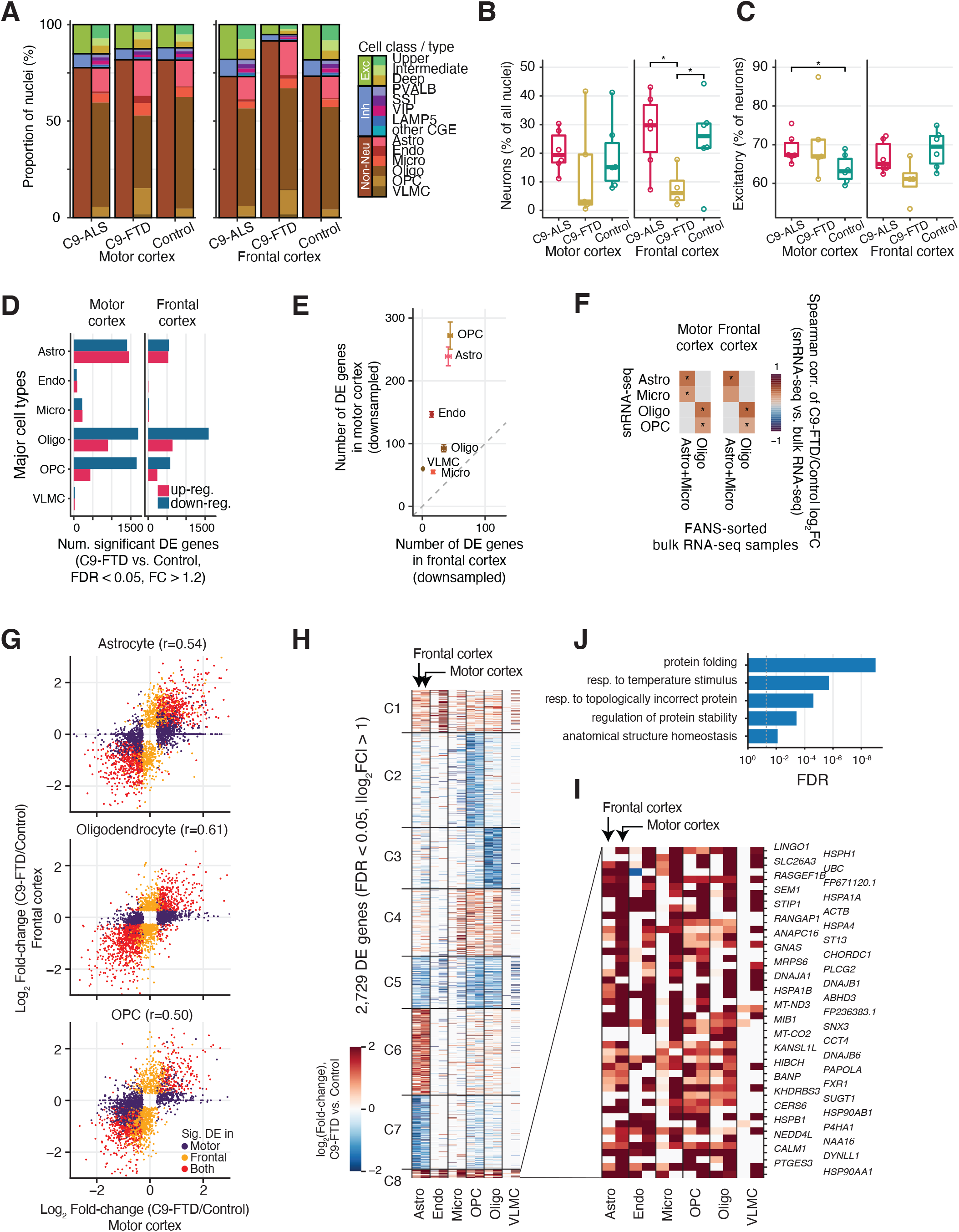
Loss of neurons and alteration of non-neuronal transcription in C9-FTD. (A) Distribution of the abundance of three cell classes and 14 major cell types across brain regions and diagnoses. (B-C) Relative abundance of neurons (B), and percent of excitatory neurons among all neurons (C). Circles represent individual donors. *, t-test p < 0.05. (D) The number of significant DE genes in FTD vs. control (FDR < 0.05, FC > 1.2) in glia. (E) Comparison of the number of FTD vs. control significant DE genes between the motor and frontal cortex after downsampling to 30 nuclei per donor in each cell type. Random downsampling was performed 10 times, and the dots and error bars represent mean ± SEM. (F) Spearman correlation between gene expression FC from snRNA-seq vs. FANS-sorted bulk RNA-seq. Correlation was performed using significant DE genes in C9-FTD vs. control identified in snRNA (FDR < 0.05, FC > 1.2). *, Spearman correlation test, FDR < 0.05. (G) Transcriptional changes in C9-FTD were consistent in two cortical regions. r, Pearson correlation coefficient. (H) K-mean clustering analysis of strongly DE genes in C9-FTD vs. control (>2 fold-change). (I) Zoom-in view of genes in cluster 8 in (H). (J) Top enriched GO terms for genes in cluster 8 (FDR<0.05). See Table S10 for the full list of GO enrichment results.

The number of high-quality neurons sampled from C9-FTD patients was too low for statistically meaningful analysis of the transcriptome. However, our data enabled high-resolution analysis of the molecular dysregulation in C9-FTD glial cells. We found thousands of DE genes in astrocytes, oligodendrocytes and OPCs, as well as a smaller but significant number of DE genes in microglia (**Fig. 6D; Table S8-9**). After downsampling to equalize the statistical power across non-neuronal cell types and brain regions, we found that astrocytes and OPCs were the most strongly affected cell types (**Fig. 6E**). We were surprised to find more DE genes in glial cells in motor cortex than in frontal cortex, in contrast to the more pronounced loss of neurons in frontal cortex (**Fig. 6E**). This observation could indicate that glial responses to neuronal degeneration and loss in C9-FTD are distributed through multiple cortical regions. In both motor and frontal cortices, we found strongly correlated differential gene expression in the single nucleus and FANS RNA-seq data for corresponding glial cell types (**Fig. 6F**). As in C9-ALS, we found that glial DE genes in C9-FTD had consistent effect sizes in both cortical regions (**Fig. 6G**).

We clustered the glial DE genes in C9-FTD, identifying groups of genes specifically affected in oligodendrocytes and OPCs (C2-C5) or in astrocytes (C6, C7) (**Fig. 6H**). These genes exhibited generally consistent effects in both motor and frontal cortices. In addition, we found a small cluster of 46 genes (C8) which were strongly upregulated across all non-neuronal cells (**Fig. 6I**). This cluster was specifically affected in motor cortex. Functional enrichment analysis showed that this group of genes was strongly associated with protein folding and regulation of protein stability (**Fig. 6J; Table S10**).

We next directly compared the disease effects in glia cells between C9-ALS and C9-FTD. To avoid double dipping, we split the 6 control donors into two groups and ran a second differential test for each glial cell type (**Methods**). We detected only a small number of shared DE genes between the two diseases (**Fig. S12A**). Also, the most strongly affected DE genes (FC > 2) in one disease showed inconsistent disease-associated changes in the other disease (**Fig. S12B-C**). These results suggest a distinct pattern of transcriptome dysregulation in C9-ALS and C9-FTD.

## Discussion

Both sporadic and C9-associated ALS and FTD are widely considered to be initiated by cell-autonomous degenerative processes in motor or von Economo neurons, respectively. Yet, mounting evidence indicates that the broader population of neuronal and non-neuronal cells play key roles in the pathology of these diseases.

Here, we used single nucleus transcriptome and epigenome sequencing from autopsied human motor and frontal cortices of cases and controls to identify the specific cell types and the biological pathways that are altered in C9-ALS or C9-FTD. These data, together with bulk RNA-seq and epigenetic (H3K27ac ChIP-seq) assays in FANS-separated nuclei, revealed distinct molecular pathologies in C9-ALS and C9-FTD across cell types and brain regions.

Previous studies have implicated all major classes of glial cells and blood-borne immune cells in the pathology of ALS (*22, 75*–*78*). Yet, it remains unclear how these findings, largely based on animal or *in vitro* models of (C9)-ALS, may translate to human ALS patients. Our analysis showed that among non-neuronal cells, astrocytes were the most strongly affected in C9-ALS. We found comparatively fewer gene expression changes in microglia and oligodendrocytes. This finding may be partly explained by the fact that our data reflect the end-of-life transcriptional disruption in C9-ALS, rather than gene expression changes at earlier stages of the disease. We detected transcriptional upregulation of markers of astrocyte activation, and of cytoskeleton and cell-surface protein remodeling in C9-ALS astrocytes. This is consistent with the hypothesis of a common astrocytic response to many forms of injury and neurodegenerative diseases, termed ‘astrogliosis’ or ‘astrocyte reactivity’, which are often accompanied by profound structural changes, including changes in different filament systems and the actin cytoskeleton (*30*). We also detected altered expression of several genes, including *ATP2B4, RANBP3L, GOLGB1, MAOAB*, that suggest dysregulation of specific cellular pathways in C9-ALS astrocytes which, if confirmed in independent studies, could provide targets for new treatment strategies.

Among the upregulated genes in C9-ALS astrocytes, a notable example is *MAOB*. MAOB protein was previously shown to be upregulated in the brains of AD patients and in the mouse models of AD (*46*). Moreover, MAOB is involved in GABA production in astrocytes, and reactive astrocytes produce and release abundant amounts of GABA as a result of the increased level of MAOB (*46*–*48*). Our findings suggest a role of astrocytic MAOB in tonic inhibition of C9-ALS neurons, which could be interpreted as a compensatory mechanism to offset glutamate-associated excitotoxicity. The latter has long been suspected as a mediator of ALS progress (*50*). Similar to C9-ALS, C9-FTD astrocytes also exhibited upregulation of activation markers (*GFAP, CD44*) as well as alterations of *ATP2B2, GOLGIB1*, and *MAOB*. Notably, C9-FTD glial cells (including astrocytes) showed upregulation of genes associated with proteostasis, which we did not detect in C9-ALS astrocytes, highlighting differences between these two diseases.

ALS causes degeneration of the giant upper motor neurons (Betz cells) located in layer 5b of the primary motor cortex, and deep layer (L5/6) excitatory neurons are generally considered vulnerable in disease (*21*). Although the Betz cells are too rare to be reliably identified in our snRNA-seq data, we found transcriptional dysregulation of L5/6 excitatory neurons which was more extensive in motor than frontal cortex. Surprisingly, we found even more extensive gene dysregulation in C9-ALS in upper layer (L2/3) excitatory neurons, which were equally afflicted in both cortical regions. This finding is intriguing given that L2/3 neurons, which mainly project intratelencephalically (IT) to connect to distant cortical regions and striatum, are greatly expanded as a proportion of the cortical neuron population in human compared with marmoset or mouse (*24, 79*). The extensive pathological alterations in L2/3 neurons in both motor and frontal cortices could thus contribute to the cognitive and behavioral symptoms in C9-ALS (*80*).

The altered gene expression in C9-ALS excitatory neurons affected biological pathways including *mitochondrial function, proteostasis*, and *protein synthesis*. Impaired mitochondrial energy production and functions (e.g., maintenance of Ca2+ homeostasis) have been described in many animal and *in vitro* models of ALS, albeit with conflicting results (*81*). Our data showed upregulation of the nucleus-encoded mitochondrial genes in deep and upper layer excitatory neurons in both cortical regions. By contrast, recent electron microscopy observations in autopsied motor cortex show decreased mitochondrial density and length in ALS patients (*82*). Mitochondria were also decreased in animal and cell models of ALS (*83*). Our findings suggest that upregulation of mitochondrial genes in C9-ALS could reflect adaptation in the surviving mitochondria to satisfy the energy demands of the neuronal cells. Indeed, neuronal energy consumption may be substantially higher in C9-ALS than healthy brains (*84*). In particular, the accumulation of DPRs and TDP43 protein aggregates or their pathogenic oligomers in C9-ALS neurons could trigger upregulation of multiple genes which encode proteins that maintain cellular homeostasis. In turn, the high energy cost of homeostasis might stimulate energy production such as mitochondrial respiratory chain and ATP synthase complexes. Consistent with this connection, we detected upregulation of proteostasis genes that encode heat shock proteins and their co-chaperones, as well as mitochondrial genes that encode proteins involved in oxidative phosphorylation (mitochondrial respiration and ATP synthesis). In turn, high metabolic activity in neurons, which rely primarily on oxidative phosphorylation in mitochondria for energy production, leads to increased ROS generation and subsequent DNA damage (*50, 85*). Indeed, increased oxidative stress and DNA damage were previously reported in neurons derived from C9-ALS and C9-FTD patients (*86*). Consistent with these findings, we detected upregulation of neuronal *SOD1* and *PRDX1* that act as scavengers of ROS (*50*), as well as upregulation of DNA damage response genes in excitatory neurons.

We also observed increased expression of genes related to translation and ribosomal function (RPL/S and MPRL/S gene families) in C9-ALS excitatory neurons. Some of the C9-ALS and C9-FTD-associated DPR proteins (poly(PR) and poly(GR)) bind to ribosomal proteins and translation elongation factors, and impair global translation (*87*–*89*). Specifically, these DPRs bind to the ribosomal tunnel and block translation, potentially explaining their toxicity in C9-ALS and C9-FTD (*90*). We therefore hypothesize that the broad upregulation of gene encoding translation system components in C9-ALS excitatory neurons represents a compensatory response to this impairment of translation.

Genes downregulated in C9-ALS excitatory neurons showed the most significant enrichments for categories associated with neuron-specific functions, reflecting neuronal degeneration associated with the disease. Of special note is *ATP2B4*, which was downregulated in excitatory neurons but upregulated in astrocytes (see above). *ATP2B4* encodes plasma membrane Ca2+-ATPase 4 (PMCA4). PMCA proteins extrude cytoplasmic Ca2+ from the neuronal cytoplasm to the extracellular milieu. Neuronal PMCAs (especially PMCA2 and 4) are extremely sensitive to glutamate-associated excitotoxicity (*91*). Downregulation of PMCA activity and protein levels were reported in aging as well as in AD and Parkinson’s diseases, and it has been suggested that these changes might further disrupt Ca2+ homeostasis, thereby promoting neuronal death (*91*). Future therapies for C9-ALS may consider stabilizing or restoring ATP2B4 expression and function to prevent neuronal degeneration.

Our epigenomic data, including single nucleus open chromatin (snATAC-seq) and histone modification H3K27ac chromatin immunoprecipitation (ChIP-seq), complement our cell type-specific transcriptome findings. The high quality of our epigenomic data from multiple donors allowed us to identify significant C9-ALS disease-associated changes of H3K27ac at promoters and distal regulatory elements, which correlated with differential gene expression and chromatin accessibility in specific glial cell types. We further found coordinated effects in C9-ALS on a large number of distal regions marked by open chromatin and H3K27ac. Notably, epigenomic alterations were more pronounced in glial cells compared with neurons, although this could partly reflect the difficulty of distinguishing cells from diverse neuronal types in sparse snATAC-seq data.

Collectively, our findings are consistent with a convergence of molecular pathology in C9-ALS excitatory neurons on protein misfolding, which triggers mitochondrial dysfunction and impairment of translation as common endpoints associated with neurodegeneration. Previous studies identified similar pathways, but the use of a variety of animal and *in vitro* models of ALS resulted in highly variable outcomes (*7, 38, 92*). Notably, at least for gene expression, we observed global upregulation of genes associated with proteostasis, oxidative phosphorylation, and translation in upper and deep layer excitatory neurons, which likely reflect compensatory mechanisms to maintain cellular homeostasis. We also detected downregulation of genes associated with multiple neuron-specific functions, apparently, reflecting the ultimate failure of these compensatory responses. Among glial cells, astrocytes showed the most extensive transcriptional dysregulation in C9-ALS and C9-FTD. Importantly, this dysregulation appears to reflect adaptive, compensatory processes, likely occurring subsequent to the degeneration of the neuronal neighbors of the astrocytes. These findings are consistent with recent genetic evidence, which suggests cell-autonomous initiation in ALS (including C9-ALS) specifically in glutamatergic neurons(*74*).

Our study provides a high-resolution view of the transcriptional and epigenetic alterations in individual cell types from the brains of C9-ALS and C9-FTD donors and offers a valuable resource for the scientific community. A limitation of this approach is that we cannot distinguish cell-type-specific alterations that represent direct effects of C9 mutation from end-of-life consequences of neurodegeneration and the resulting disease which are present in the C9-ALS or C9-FTD cases. Also, our methods do not allow for a spatial resolution of the ALS-associated transcriptome dysregulation, which provides valuable information of how this dysregulation participates in the spread of ALS or FTD pathology (*93*). Although we confirmed that differential expression of some genes results in the corresponding changes in protein expression, it is possible that some disease-associated mRNA effects are counteracted by homeostatic regulation of translation, justifying future large-scale proteomic studies of ALS and FTD brain. Lastly, the pronounced loss of neurons in C9-FTD patients precluded statistically meaningful analysis of the neuronal transcriptome. Overall, this work demonstrates the value of large-scale single cell/nucleus studies in patients to establish the cell-type-specific molecular pathology of C9-ALS and C9-FTD, which will be essential for developing targeted disease-modifying therapies.

## Materials and Methods

### Experimental Design

This study was designed to determine transcriptional and epigenetic alterations in the brains of ALS and FTD donors with the repeat expansion in *C9orf72* (C9) gene in single cell resolution. For this purpose, we performed snRNA-seq and snATAC-seq in postmortem samples from C9-ALS (N=6), C9-FTD (N=5) and pathologically normal donors (N=6). To compare the impact across different cortical regions, these studies were performed in motor cortex and frontal cortex, which have been considered to be afflicted the most in ALS and FTD, respectively. For validation, the single nucleus studies were complemented by RNA-seq and epigenetic (H3K27ac ChIP-seq) assays in FANS-separated nuclei from major brain cell types, as well as by automated Western blotting in bulk motor cortex tissue. No statistical methods were used to predetermine sample size because effect sizes were unknown before experiments.

### Nuclei isolation and snRNA/snATAC-seq on the 10X Genomics platform

Nuclei were isolated from 50-70 mg frozen brain tissue (**Table S1**) mainly as described (*94*) (see details in **Suppl. Methods**). Single-nucleus capture and library preparation were then performed using the 10X Genomics Single-Cell 3’ protocols without any modifications. Matched control, C9-ALS and C9-FTD samples were loaded on the same 10X chip to minimize potential batch effects. Single-nucleus libraries from individual samples were pooled and sequenced on the NovaSeq 6000 machine with an average depth of ∼10,870 unique molecular identifiers (UMIs) per nucleus for snRNA-seq and ∼8,223 unique fragments per nucleus for snATAC-seq.

### Fluorescence-activated nuclei sorting (FANS) and nuclear RNA-seq of the three populations of the human brain cells (neurons, oligodendrocyte lineage cells, and other glia)

The isolation of brain nuclei prior to the flow cytometry separation was performed mainly as described (*95*) (**Suppl. Methods**). FANS was then used to isolate neurons (NeuN+SOX10-population), oligodendrocyte lineage cells (NeuN-SOX10+ population), consisting of mature oligodendrocytes and a smaller population of OPCs, and a third population, “other glia” (NeuN-SOX10-) that consisted mainly of astrocytes and microglia. From each sample (**Table S1**), 150-300 thousand (K) nuclei of each cell type were collected. RNA was then extracted from these nuclei, and 10 ng of the RNA preparations were used to prepare RNA-seq libraries using the SMARTer Stranded Total RNA-seq Kit, Pico-Input v2 (Takara). Libraries were sequenced on a NovaSeq 6000 (Illumina), using paired-end (PE) 100 cycles protocol to an average of 120 million read pairs per sample.

### FANS and H3K27ac ChIP-seq of the four populations of the brain cells (neurons, oligodendrocytes, astrocytes, and microglia)

FANS to isolate neurons, oligodendrocytes, astrocytes, and microglia was performed mainly as described in the previous section with modifications, including adding antibodies against interferon response factor 5 (IRF5) (**Suppl. Methods**). From each sample (**Table S1)**, 100-150K nuclei of each cell type were collected and used in a native ChIP assay (Suppl. Methods) (*96*). ChIP-Seq libraries were prepared with the NEBNext Ultra II DNA Library Prep Kit for Illumina (New England Biolabs) and sequenced on a NovaSeq 6000, using PE100 protocol to an average of 60 million read pairs per sample. For each cell type and diagnostic group, three input control samples obtained from MNAse-digested DNA were prepared and sequenced.

### Jess/Wes automated, multiplex Western blot assay

Capillary Western analyses were performed in the motor cortex samples from C9-ALS and control donors (**Table S1**) using the ProteinSimple Jess-Wes System (San Jose, CA, USA) (**Suppl. Methods**). A two-sided Welch’s t-test was used to compare the immunoreactive signals between C9-ALS and control, where the averaged signals between two replicates for each donor were used as observations (6 donors each for C9-ALS and control), and the comparisons with a p-value less than 0.05 were deemed significant.

### GFAP immunofluorescence

The assay was performed on frozen human postmortem motor cortex blocks from control and C9-ALS samples (**Suppl. Methods**). Double labeling (GFAP and DAPI) was confirmed by acquiring z-stacks at 1 to 0.5 μm intervals through cells of interest and by maximum intensity projection (MIP) views as described (*97*).

### snRNA-seq data processing

snRNA-seq reads were mapped to the human GRCh38 genome using 10x Genomics Cell Ranger (*98*). CellBender (*99*) was used to distinguish cell-containing from cell-free droplets and retrieve noise-free cell-by-gene quantification tables. Scublet (*100*) was used to compute the doublet score for each nucleus. Nuclei with fewer than 500 detected genes, more than 1% reads that mapped to the mitochondrial genome, or a doublet score greater than 0.2 were excluded from further analysis. Seurat (*101*) was used to perform Sctransform normalization (*102*), Principal component analysis (PCA), Harmony integration (*103*), Leiden clustering, and UMAP (Uniform Manifold Approximation and Projection) (*104*) visualization. Clusters were annotated using the expression of selected marked genes (**Fig. S3B-D**). Seurat’s label transfer functions were used to estimate how well our annotation matched with the cell types annotated in the recent human motor cortex single nuclei study by the Allen Institute for Brain Science (*24*). The model-based analysis of single-cell transcriptomes (MAST) (*28*) was used to perform differential expression analysis, systematically accounting for predefined covariates. GO enrichment analysis was done using WebGestalt (*34*). See **Suppl. Methods** for details.

### Bulk FANS-sorted nuclear RNA-seq data processing

FastQC (*105*) was used to examine the quality of the RNA-seq reads. Reads were trimmed using Trim Galore (a wrapper tool powered by Cutadapt (*106*)) in paired-end mode. Trimmed reads were then mapped to the human hg38 genome and the GENCODE annotated transcriptome (release V35) with STAR (Spliced Transcripts Alignment to a Reference) (*107*). Gene expression was estimated using RSEM (RNA-Seq by Expectation Maximization (*108*). edgeR (*109*) was used to perform differential expression tests separately for each sorted population in each brain region.

### snATAC-seq data processing

snATAC-seq reads were mapped to the human GRCh38 genome using 10x Genomics Cell Ranger ATAC (*110*). ArchR (*111*) was used in the downstream analysis. Fragment size distribution in each sample was inspected for nucleosomal periodicity, and nuclei with TSS (Transcription Start Site) enrichment score less than 4 and a number of unique fragments less than 1000 were removed. After doublet inference and removal, a cell-by-tile matrix containing insertion counts across genome-wide 500-bp bins was created. Dimensionality reduction with iterative Latent Semantic Indexing (LSI), Louvain clustering and UMAP visualization were then performed. Gene activity scores were computed using the model implemented in ArchR, accounting for both the accessibility within the entire gene body and the activity of putative distal regulatory elements. Clusters were annotated using labels transferred from our snRNA-seq dataset. Reproducible peaks were called using MACS2 (*112*). Differential chromatin accessibility was assessed with the Wilcoxon test in both gene-based levels (with gene activity scores) and peak-based levels. ChromVAR (*72*) was used to estimate chromatin accessibility within peaks sharing the same transcription factor (TF) motif while controlling for technical biases. See **Suppl. Methods** for details.

### H3K27ac ChIP-seq processing

Trimmed reads were mapped to the human hg38 genome BWA-MEM2 (*113*) and filtered to remove multi-mapping reads and low-quality alignments using Samtools (*114*). Reads mapping to ENCODE-blacklisted genomic regions were excluded using BEDTools (*115*). H3K27ac-enriched peaks were detected using MACS2, including input controls for each cell type and condition, as previously described (*116*). Promoter H3K27ac signal was computed using the from deepTools (*117*). Differential peaks between C9-ALS and control for each FANS-sort population were identified using DiffBind (*118*) in edgeR (*109*) mode.

### Statistical analysis

All statistical analyses were performed in R or Python. The specific statistical test used for each figure is included in the figure legend, main text and the detailed materials and methods section in the supplementary materials. Benjamini and Hochberg’s false discovery rate (FDR) method was used to adjust for multiple comparison, and FDR < 0.05 was used as cutoff to call significant except where stated otherwise.

## List of supplementary materials

Detailed Materials and Methods.

### Supplementary Figures

- Fig. S1. Quality control (QC) metrics for snRNA-seq data.
- Fig. S2. The clusters identified in snRNA-seq were not biased by known covariates.
- Fig. S3. Annotations of the identified cellular populations in snRNA-seq based on the expression of known markers.
- Fig. S4. Comparison of the cell types identified in our snRNA-seq dataset with a previously published dataset.
- Fig. S5. Fluorescence-activated nuclei sorting (FANS) isolation of nuclei of major brain cell types.
- Fig. S6. Quantification of proteins encoded by genes dysregulated in C9-ALS astrocytes using automated Western blot analysis.
- Fig. S7. Quantification of proteins encoded by genes dysregulated in C9-ALS excitatory neurons using automated Western blot analysis.
- Fig. S8. Functional enrichment analysis of differentially expressed (DE) genes in upper-and deep-layer excitatory neurons.
- Fig. S9. QC metrics used in the snATAC-seq data pre-processing.
- Fig. S10. The epigenetic landscape of C9-ALS brain cells determined by snATAC-seq.
- Fig. S11. ChromVAR scores in non-redundant transcription factor (TF) motif archetypes.
- Fig. S12. Comparison of the disease effect on gene expression in non-neuronal cells in C9-ALS and C9-FTD.

Supplementary Tables

- Table S1. Metadata of the samples used in this study.
- Table S2. Metadata and cell type annotations of all high-quality nuclei in our snRNA-seq dataset.
- Table S3. List of C9-ALS vs. control differentially expressed genes identified in each major cell type in motor cortex.
- Table S4. List of C9-ALS vs. control differentially expressed genes identified in each major cell type in frontal cortex.
- Table S5. Gene Ontology enrichment results for C9-ALS vs. control differentially expressed genes.
- Table S6. Metadata and cell type annotations of all high-quality nuclei in our snATAC-seq dataset.
- Table S7. H3K27ac differential peaks between C9-ALS and control identified in ChIP-seq.
- Table S8. List of C9-FTD vs. control differentially expressed genes identified in each non-neuronal cell type in motor cortex.
- Table S9. List of C9-FTD vs. control differentially expressed genes identified in each non-neuronal cell type in frontal cortex.
- Table S10. Gene Ontology enrichment results for genes upregulated across all non-neuronal cell types (Cluster 8) in C9-FTD vs. control.

References (119–129) (references only cited in SM)

## Acknowledgements

We gratefully acknowledge Eugene Koonin for helpful comments. This work was supported by VA Merit Awards BX003625 and BX005585 (S.D.); PsychEncode consortium NIH/National Institute of Mental Health U01MH122590 (S.D.) and U01MH122592 (E.A.M.); R01AG067151 (V.B.B). All authors declare that they have no conflicts of interest. All sequencing datasets have been deposited in the Gene Expression Omnibus repository with the accession number GSExxxxxx. Codes used in this study are available at https://github.com/hoholee/C9_ALS_FTD_single_nuclei_transcriptome_epigenome.

## Author contributions

Conceptualization: JL, VVB, EAM, SD;

Methodology: JL, MKJ, JC, AK, PZ, EAM, SD;

Software: JL, JC, EAM;

Validation: MKJ;

Formal analysis: JL, JC, EAM;

Investigation: JL, MKJ, JC, AK, PZ, EAM,

SD Resources: MG, LJP, EE-C, DWD, VVB, SD;

Data Curation: JL, JC, AK, VVB, SD;

Writing – original draft: JL, MKJ, JC, AK, EAM, SD;

Writing – review & editing: JL, MKJ, JF, AK, VVB, EAM, SD;

Visualization: JL, JC, EAM;

Supervision: EAM, SD;

Project administration: EAM, SD;

Funding acquisition: EAM, SD.

