## Supplementary Materials for "Divergent impacts of *C9orf72* repeat expansion on neurons and glia in ALS and FTD"

#### Detailed Materials and Methods

##### Human samples

Human frozen post-mortem tissues were obtained from the Mayo Clinic Brain Bank (Jacksonville, FL USA). Data analysis was conducted as exempt human research, considering that these were postmortem brain samples, and they were not specifically collected for this study. The samples were dissected from the dorsolateral prefrontal cortex (Brodmann area 9; hereafter “frontal cortex”) and/or motor cortex (Brodmann area 4). A total of 17 donors with C9-ALS (N=6), C9-FTD (N=5), or found pathologically normal (N=6) were used for snRNA-seq, snATAC-seq, bulk RNA-seq in FANS-separated nuclei from major brain cell type, and automated Western blotting studies (**Table S1**). A separate group of C9-ALS (N=6) and pathologically normal (N=9) donors was used in H3K27ac ChIP-seq studies **(Table S1)**. All subjects were confirmed negative for protein-coding mutations in *TARDBP, FUS, NEK1, GRN, MAPT,* or *TBK1*. In all C9-ALS and C9-FTD samples, TDP-43 pathology was confirmed based on current consensus criteria, which investigates the cortical and subcortical distribution of TDP-43 neuropathologic inclusions (*119*, *120*). *C9orf72* repeat expansions were confirmed via repeat-primed PCR and Southern blotting.

##### Nuclei isolation and snRNA/snATAC-seq using the 10X Genomics platform

Nuclei were isolated from 50-70 mg frozen brain tissue mostly as described (*94*). For each donor, brain tissues from the motor cortex and frontal cortex were dissected with a scalpel blade. The tissues were then homogenized on ice in 5mL lysis buffer (0.32 M sucrose, 3 mM $Mg(Ac)_2$, 3 mM $CaCl_2$, 0.1 mM EDTA, 1 mM DTT, 0.1% Triton-X, and 10 mM Tris-HCl pH 8.0 in DEPC-treated water), containing 0.4 U/µl freshly added Recombinant RNase Inhibitor (RRI; Takara, Cat. # 2313A) until no chunks of tissue were visible in the tissue suspension (~20 strokes) using a glass Dounce homogenizer (Thomas Scientific; Cat. # 3431D76; size A). The homogenized tissues were filtered through a 40 µm strainer (Thermofisher; Cat. # 22-363-547) and transferred to a 30 mL thick polycarbonate ultracentrifuge tube (Beckman Coulter; Cat. # 355631). Nine mL of the sucrose buffer (1.8 M Sucrose, 3 mM $Mg(Ac)_2$, 1 mM DTT, and 10 mM Tris-HCl pH 8.0 in DEPC-treated water) were added to the bottom of the tubes with the tissue homogenate, and the tubes were centrifuged at 34,400 RPM (107,164 FCF) for 2.5 hours at 4°C, using a swing bucket rotor (SW28). The supernatant was aspirated, and the nuclear pellet was submerged on ice for 20 min in 250 uL of DEPC-treated water-based PBS, containing 1% BSA and 0.4 U/µL RRI. The nuclear pellet was then resuspended and filtered twice through a 30 µm cell strainer (Miltenyi Pre-separation Filters; Cat. # 130-041-407). Nuclei were counted using a hemocytometer and diluted to ~2,000 nuclei/µL before performing single-nucleus capture on the 10X Genomics Single-Cell 3’ system. The 10X capture and library preparation protocols were used without modification. Matched control, C9-ALS and C9-FTD samples were loaded on the same 10X chip to minimize potential batch effects. Single-nucleus libraries from individual samples were pooled and sequenced on the NovaSeq 6000 instrument (Illumina) with an average depth of ~10,870 unique molecular identifiers (UMIs) per nucleus for snRNA-seq and ~8,223 unique fragments per nucleus for snATAC-seq.

##### Fluorescence-activated nuclei sorting (FANS) and nuclear RNA-seq of the three populations of the human brain cells (neurons, oligodendrocyte lineage cells, and other glia)

The isolation of brain nuclei prior to the flow cytometry separation was performed mostly as described (*95*). Tissue (~300 mg) was homogenized in ice-cold lysis buffer (320 mM sucrose, 5 mM $CaCl_2$, 3 mM $Mg(Ac)_2$, 0.1 mM EDTA, 0.1% Triton X-100, 1 mM DTT, 1 U/µl RRI, and 10mM Tris-HCl, pH 8.0), underlaid with the sucrose buffer (1.8 M Sucrose, 3 mM $Mg(Ac)_2$, 1 mM DTT, 0.4 U/µl RRI, and 10 mM Tris-HCl pH 8.0), and centrifuged for 1h at 24,000 rpm. The nuclear pellets were then resuspended in the antibody-incubation buffer (0.5% BSA, 3mM $MgCl_2$, 1 U/µl RRI, and 10 mM Tris-HCl, pH 8.0) and incubated with antibodies against NeuN and SOX10 for 1 hr at 4°C. FANS method was then used to isolate neurons (NeuN+SOX10- population), oligodendrocyte lineage cells (NeuN-SOX10+ population), consisting of mature oligodendrocytes and a smaller population of

OPCs, and a third population (NeuN-SOX10-; hereafter "other glia") that mostly consisted of astrocytes and microglia.

NeuN (also known as RNA-Binding Protein RBFOX3) is a well-established marker of neuronal nuclei (*121*). SOX10 is a transcription factor specifically expressed in oligodendrocyte lineage cells. The application of anti-SOX10 antibodies to isolate oligodendrocyte lineage nuclei was described by Frisen and colleagues (*122*). We used Alexa488-conjugated anti-NeuN antibodies (1:1000 dilution, Millipore, Cat. # MAB377x) and anti-SOX10 antibodies (R&D Systems, Cat. # AF2864) that were custom conjugated to Alexa647 (1:150 dilution, Cat# FCMAB317PE, Millipore). DNA stain DAPI was used to label intact nuclei (**see Fig. S5 for details**). From each sample, we collected 250-300 thousand (K) nuclei of neurons, 250-300K of oligodendrocyte lineage cells, and 150-200K nuclei of other glia. Nuclei were collected directly to the lysis buffer from PicoPure RNA Isolation Kit (ThermoFisher Scientific, Cat. # KIT024), which was then used for RNA isolation. RNA-seq libraries were prepared with the SMARTer Stranded Total RNA-seq Kit, Pico-Input v2 (Takara, Cat. # 634414) from 10ng of the RNA. Libraries were sequenced on a NovaSeq 6000 instrument (Illumina), using paired-end 100 cycles protocol to an average of 120 million read pairs per sample.

**FANS and H3K27ac ChIP-seq of the four populations of the brain cells (neurons, oligodendrocyte lineage, astrocytes, and microglia)**

FANS protocol to isolate neurons, oligodendrocyte lineage, astrocytes, and microglia was performed as described in the previous section, except removing RRI from all buffers, adding 0.1 mM benzamidine, 0.1 mM phenylmethylsulfonyl fluoride (PMSF) to the lysis buffer, and adding antibodies against interferon response factor 5 (IRF5). IRF5 is highly enriched in cells of myeloid origin, including microglia (*123*). The IRF5 antibodies allowed the separation of astrocytes and microglia within the other glia (NeuN-OLIG10-) population (**see Fig. S5 for details**). In this FANS protocol, we used PE-conjugated anti-NeuN antibodies (1:1,000 dilution, Millipore, Cat. # FCMAB317PE), anti-SOX10 antibodies (R&D Systems, Cat. # AF2864), which were custom-conjugated to Alexa647 (1:150 dilution), and Alexa488-conjugated anti-IRF5 antibodies (1:200 dilution, R&D Systems, Cat. # IC4508G).

We employed the native ChIP protocol (N-ChIP) in which chromatin fragmentation is performed using micrococcal nuclease (MNase) without crosslinking proteins to DNA (*96*). 100-150K of each cell type were collected and used for each ChIP reaction. We used anti-H3K27ac antibodies from Active Motif (Cat# 39133; rabbit polyclonal, 3 µg per sample). ChIP-Seq libraries were prepared with the NEBNext Ultra II DNA Library Prep Kit for Illumina (New England Biolabs, Cat. # E7645). The resulting libraries were sequenced on a NovaSeq 6000 instrument (Illumina), using paired-end 100 cycles protocol, to an average of 60 million read pairs per sample. For each diagnostic group, three input control samples obtained from MNAse-digested DNA were prepared and sequenced.

**Jess/Wes automated, multiplex Western blot assay**

Capillary Western analyses were performed using the ProteinSimple Jess-Wes System (San Jose, CA, USA). The following reagents were used: Jess/Wes-Separation Module [2-40kDa Kit (Cat. # SM-W012) 12-230 kDa Kit (Cat. # SM-W004) and 66-440 kDa kit (Cat. # SM-W007)]; Jess/Wes- anti-rabbit, anti-mouse, and anti-goat Detection Module kit (Cat. # DM-001, DM-002 and DM-006, respectively); anti-mouse, anti-rabbit and anti-goat secondary antibodies (Cat. # 042-205, 042-206 and 043-522-2, respectively); antibody diluent (Cat # 042-203); EZ standard Pack 1, 3 and 5, containing biotinylated ladder (molecular weight 12-230 kDa, 66-440 kDa, and 2-40 kDa, respectively); fluorescent (FL) standards, containing 29 kDa (Cat. # PS-ST01EZ), 90 kDa (PS-ST03EZ), and 26 kDa (PS-ST05EZ) system controls; dithiothreitol (DTT); streptavidin-HRP; luminol-S, peroxide; sample buffer (Cat. # 042-195); and wash buffer (Cat. # 042-202). The Separation Module kit included capillary cartridge and pre-filled microplates. Primary Antibodies used were specific for HSP90 (Mouse, R&D Systems Cat, # MAB3286), GFAP (Mouse, Sigma Cat, # G3893), CD44 (Mouse, R&D Systems, Cat. # BBA10), CHI3L1 (Goat, R&D Systems, Cat. # AF2599), HSP70 (Rabbit, R&D Systems, Cat. # AF1663), HSP27 (Rabbit, R&D Systems, Cat # AF1580), RANBP3L (Rabbit, Novus Cat # NBP2-38347), KCND3/Kv4.3 (Rabbit, Novus Cat. # NBP2-76945), DNMT3A (Mouse, Novus Cat # NBP120-13888), NEFL (Mouse R&D Cat

### MAB22163), Clusterin (Mouse, Novus Cat. # MAB2937), RAP1GAP (Rabbit, Novus Cat. # NBP1-53072), UbiquitinB (Mouse, Novus Cat. # NBP3-07163), and TGFB2 (Goat, Novus Cat # AB-112-NA).

The assay was performed in the motor cortex samples from C9-ALS and control donors (**Table S1**). For each subject, ~ 80-100 mg of tissue were homogenized in lysis buffer (3µl/mg of tissue) [0.32M Sucrose, 5mM $CaCl_2$, 3mM $Mg(Ac)_2$, 150 mM NaCl, 0.1 mM EDTA, 0.1% Triton X100, 3% Igepal.NP-40 (v/v), 1% sodium deoxycholate (w/v), 0.1% SDS (w/v), 1mM PMSF, Protease Inhibitor Cocktail (Sigma, St. Louis, MO; 1:100 v/v), and 10mM Tris (pH 8.0)] using a ME220 focused ultrasonicator (Covaris, Cat # 500506). Burst setting of 20 Sec, power setting of 180 watts were used; the suspension was chilled at 4 °C between ultrasonic bursts. Following homogenization, an additional 0.2mM of PMSF was added to each sample, and the samples were incubated on ice for 30 min. The homogenates were then centrifuged at 10,000 g for 20 min at 4 °C, and the supernatant (whole-tissue extract) was transferred to a new tube. Total protein concentrations of the whole-tissue extracts were measured using Qubit Protein Assay (ThermoFisher Scientific, Cat # Q33211). Samples were stored at −80 °C.

On the day of an experiment, samples (whole-tissue extracts) were defrosted and diluted with 0.1X Sample Buffer (from a 10X stock). The diluted samples were then combined with 5X Fluorescent (5X FL) Master Mix (containing 5X sample buffer, 5X fluorescent standard, and 200 mM DTT) in a 4:1 ratio and denatured by heating at 95 °C for 5 min using TruTemp DNA Microheating System (Robbins Scientific, Cat # 1057-30-0). The Fluorescent Master Mix contains three fluorescent proteins that serve to normalize the separation distance within each capillary, as the molecular weight ladder is loaded only to the first capillary. Following the denaturation step, the samples, blocking reagent, chemiluminescent substrate (mixture of luminol-S and peroxide in a 1:1 ratio), washing buffer, HRP-conjugated secondary antibodies, and a primary antibody were dispensed into designated wells in the assay plate. Following plate-loading, the samples were subjected to a fully automated capillary separation electrophoresis and immunodetection. One target protein was tested in each assay, and the corresponding primary antibodies were diluted to ensure the linear dynamic range of detection for each protein. The following dilutions were used: 1:5 dilutions for clusterin (CLU), 1:10 dilution for CHI3L1, CD44, HSP90, KCND3/Kv4.3, DNMT3A, RAP1GAP, Ubiquitin B, and 1:20 dilution for GFAP, HSP27, NRFL, RANBP3L.

For the statistical analysis, ProteinSimple Compass for SW software was used to extract areas under peak values, which were used to analyze the immunoreactive signals. A two-sided Welch's t-test was used to compare the signals between C9-ALS and control, where the averaged signals between two replicates for each donor were used as observations (6 donors each for C9-ALS and control), and the comparisons with a p-value less than 0.05 were deemed significant.

**GFAP immunofluorescence**

Frozen human postmortem motor cortex blocks from control and C9-ALS samples were embedded in OCT (TissueTek Sakura) and cryosectioned at 10µm at −20 °C (Thermo Cryostar). Sections were placed onto Superfrost Plus glass slides (Fisher Scientific), dried for 20 min at −20 °C, sealed, and stored at −80 °C until use. Immunofluorescence staining was performed as described (*97*). In short, sections were washed in 1x PBS for 60 min, treated with 0.5% sodium borohydride to remove aldehydes, rinsed in PBS, incubated for 10 min in 3% hydrogen peroxide to inhibit endogenous peroxidase, and then incubated in blocking solution (1xPBS, 0.3% Triton-X 100, 10% Normal Goat Serum, 1% Bovine Serum Albumin) for 2 hours at room temperature (RT). This was followed by overnight incubation at 4 °C in blocking solution with rabbit anti-GFAP primary antibodies (1:1,000; Millipore Sigma, Cat. #AB5804). Sections were then washed 3 times in 1x PBS (10 min each), and incubated with goat anti-rabbit IgG Alexa Fluor 594 secondary antibodies (1:500, Invitrogen). Subsequently, sections were incubated in 0.1% Sudan Black in 70% ethanol for 5 min at RT to suppress autofluorescence, washed in Tris buffer saline (pH 7.6) 3 times (15 min each), cover-slipped with ProLong Gold antifade reagent with DAPI (Invitrogen), and seal with nail polish. Fluorescent labeled sections were imaged using 63x oil-immersion objectives. The images were captured on an inverted Zeiss LSM 700 confocal microscope (Zeiss, Germany). Double labeling was confirmed by acquiring z-stacks at 1 to 0.5 µm intervals through cells of interest and by maximum intensity projection (MIP) views as described (*97*). The images were acquired in 3 sections per subject and 3 fields in each section.

**snRNA-seq data preprocessing, clustering and cell type annotation**

snRNA-seq reads were mapped to the human GRCh38 genome with a custom pre-mRNA annotation (modified RefSef gene annotation where each gene transcript locus was listed as an exon) using 10x Genomics Cell Ranger (v3.0.2) (*98*) with the default parameters. The raw feature-by-droplet matrices generated from the "cellranger count" module were then fed to CellBender (v0.2.0) (*99*) to distinguish cell-containing from cell-free droplets and retrieve noise-free cell-by-gene quantification tables using the following parameters: "--expected-cells 1500 --total-droplets-included 18000 --model full --epochs 150 --cuda --low-count-threshold 15 --fpr 0.01 --learning-rate 0.0001 --posterior-batch-size 5 --cells-posterior-reg-calc 50".

Scublet (v0.2.3) (*100*) was then used in the CellBender-generated h5 file for each sample to compute the doublet score for each nucleus with an expected doublet rate of 0.06. Based on the simulated doublet histogram of doublet scores, we set the doublet score cutoff to 0.2 and this threshold worked well for all samples.

Next, Seurat (v4.0.4) (*101*) was used to create a merged object containing all the samples, and to perform the following downstream processing. Nuclei with fewer than 500 detected genes, more than 1% reads that mapped to the mitochondrial genome, or a doublet score greater than 0.2 were excluded from further analysis. Sctransform (*102*) normalization was performed on the remaining nuclei, with the mitochondrial mapping percentage and the sequencing batches set as the confounding sources of variation to be removed. Principal component analysis (PCA) was run with the top 3000 highly variable genes in the normalized nuclei-by-gene matrix, and the first 30 principal components were used to run Harmony (v0.1.0) (*103*) integration with sample IDs, donor IDs and sequencing batches set as covariants. Then, a k-nearest-neighbor graph was constructed with k set as 30, and the nuclei were clustered using Leiden clustering with a resolution of 1. This gave rise to 29 clusters, and UMAP (Uniform Manifold Approximation and Projection) (*104*) was used for visualization (Fig. 1A, the leftmost panel).

The nuclei were then classified into three classes (excitatory neurons, inhibitory neurons, and non-neurons) using several well-established marker genes (Fig. S3A). For each of these three classes, a subset of nuclei was created from the full dataset, and the aforementioned Sctransform normalization, PCA, Harmony integration and Leiden clustering steps were run on each of these subsets. This iterative clustering process reselects the highly variable genes in the context of each cell class and hence may give better resolution in finding distinct subclusters. With the Leiden resolution set as 1.5, 0.5 and 1, we identified 24, 23, and 23 subclusters for excitatory neurons, inhibitory neurons and non-neurons, respectively. These subclusters were annotated using the expression of selected marked genes (Fig. S3B-D). Note that six subclusters seemed to express marker genes of multiple cell classes (for example an excitatory subcluster with expressions of oligodendrocyte marker genes), which were potential doublets that failed to be captured by the Scublet pipeline. We labeled these subclusters as "ambiguous" and removed them from further consideration. We also noticed one excitatory neuron subcluster that highly expressed the *TUBB2A* gene but had a low number of detected genes, and one non-neuron subcluster that expressed marker genes of immune cells (but no expression of microglia marker genes). We also excluded these two subclusters from the downstream analysis. After annotation, we identified 49 distinct subpopulations of excitatory neurons, inhibitory neurons and non-neuronal cells. To estimate how well our annotation matched with the cell types annotated in the recent human motor cortex single nuclei study by the Allen Institute for Brain Science (*24*), we used Seurat's "FindTransferAnchors" and "TransferData" functions to transfer the annotation labels from the AIBS reference dataset to our dataset. A confusion matrix was constructed with the prediction labels (label with the highest prediction score) from the data transfer and the labels from our annotation, and the adjusted rand index was calculated to measure the similarity between the two (Fig. S4). Cell type nomenclature mostly followed the common cell type nomenclature (CCN) convention presented by the Allen Institute for Brain Science (*24*, *124*), with slight modifications to reflect the specific marker genes found in our dataset.

**Differential expression analysis in snRNA-seq**

To ensure a statistically meaningful number of cells from each donor for the differential expression analysis, we grouped the subpopulations into 14 major cell types with shared marker genes. In these major cell types, there were at least 15 high-quality nuclei in at least 4 out of the 6 donors in each group (the combination of diagnosis and brain region). For the comparison between C9-FTD and control samples, we focused on the non-neuronal cell types as the C9-FTD samples showed depletion in the numbers of neuronal nuclei (Fig. 5A). To identify the differentially expressed genes (DEGs) between disease (C9-ALS or C9-FTD) and control across these major cell types in each brain region, we used the model-based analysis of single-cell transcriptomes (MAST, v1.18.0) (*28*), where a mixed-effect hurdle model was employed to model the snRNA expression data as a mixture of a binomial and normal distribution while systematically accounting for predefined covariates. For each comparison, the raw counts of chosen nuclei in question were extracted from the "RNA" slot of the Seurat R object and then normalized to $log_2$CPM (counts per million). Genes that were expressed in at least 10% of the selected nuclei were kept. The following linear mixed model was then fit with MAST:

$$y \sim D + G + U + M + A + S + B + (1|I)$$

Here, $y$ is the $log_2$-normalized count of the gene; $D$ is the diagnosis (ALS, FTD or control); $G$ is the number of genes detected, $U$ is the number of UMIs; $M$ is the percentage of reads mapped to mitochondrial genes; $A$ is the age of the donor; $S$ is the sex of the donor; $B$ is the sequencing batch of the sample; and $I$ is the ID of the donor. $G$, $U$, $M$ and $A$ were centered and scaled across the selected nuclei in comparison. The donor ID $I$ was modeled as a random-effect term while the rest of the covariates were modeled as fixed-effect terms.

Next, a likelihood ratio test (LRT) was performed to identify DEGs by comparing the model with and without the diagnosis term. Hurdle p-values were reported by MAST, and Benjamini & Hochberg's false discovery rate (FDR) method was used to adjust p-values for multiple comparisons. The MAST model also reported fold-changes (FC) due to the disease effect (hereafter referred to as model FC), which were contributed by both the continuous component (nonzero expression) and the discrete component (expressed or not) of the hurdle model. We also computed more straightforward fold changes between the groups (hereafter referred to as average FC) by subtracting the mean $log_2$CPM of selected nuclei in the control sample from the mean $log_2$CPM of nuclei in the disease sample. In most cases these two fold-changes were concordant, and we excluded genes with extremely high model FC that deviated from the average FC. Genes were defined as significantly differentially expressed if they passed all of the following criteria: the FDR was less than 0.05; the absolute model FC was greater than 20%; both the continuous and the discrete fits of the hurdle model were convergent; the upper and lower bounds of the 95% confidence model had the same sign; the difference between the $log_2$(model FC) and $log_2$(average FC) was less than 2.

To control for the number of nuclei when comparing the disease effect across the major cell types, we randomly downsampled the dataset to 30 nuclei per donor in each cell type, and performed the same MAST test as described above. The downsample analysis was done 10 times, and the number of significant DEGs for each cell type was reported as mean ± SEM (standard error of the mean).

To avoid double dipping when comparing the disease effect in non-neuronal cells between C9-ALS and C9-FTD, we split the 6 control donors into two groups (group 1: C1, C2 and C3, two males and one female; group 2: C4, C5 and C6, one male and two females). The same MAST test was run as described above, except that the group 1 control donors were used in the C9-ALS vs. control comparison while group 2 control donors were used in the C9-FTD vs. control comparison. The overlaps of significant DEGs between the two comparisons were reported, and the correlations of model fold-changes were estimated using the Pearson correlation coefficient.

**Gene Ontology (GO) enrichment test**

GO enrichment analysis was done using WebGestalt (*34*). Up- and Down-regulated DEGs identified in each cell type and each comparison were used separately as input, and all the expressed genes in the corresponding group were set as the reference gene set (background). GO terms in the functional databases "Biological Process noRedundant", "Cellular Component noRedundant" and "Molecular Function noRedundant" were used, and only terms with 25 to 500 genes were included in the over-representation analysis. Significantly enriched GO terms (FDR < 0.05) were then clustered by Affinity Propagation to further remove reductant terms.

To compare the effect size of differential expression between motor cortex and frontal cortex, we calculated the fold-change difference ($\Delta\ log_2FC$) between the two brain regions for genes associated with each GO term of interest. The $\Delta\ log_2FC$ for all expressed genes (labeled as "all genes" in Fig. 4D and Fig. S8A) and all DE genes (labeled as "DE genes") were also computed and used as baseline control. T-test was used to test whether the $\Delta\ log_2FC$ in each group of genes were significantly different from the $\Delta\ log_2FC$ of the "all genes" control set.

**Bulk FANS-sorted nuclear RNA-seq data processing**

FastQC (v0.11.8) (*105*) was used to examine the quality of the RNA-seq reads. Reads were then trimmed to remove sequencing adapters and low-quality sequences (minimum Phred score 20) using Trim Galore (v0.5.0, a wrapper tool powered by Cutadapt (*106*)) in paired-end mode. The first 3 bp from the 5′ end of read 1 were also removed. Trimmed reads were then mapped to the human hg38 genome and the GENCODE annotated transcriptome (release V35) with STAR (Spliced Transcripts Alignment to a Reference, v 2.7.1a) (*107*). Gene expression was estimated using RSEM (RNA-Seq by Expectation Maximization, v1.2.30) (*108*). Gene-level 'expected count' from the RSEM results were rounded and fed into edgeR (v3.34.1) (*109*) to perform differential expression tests separately for each sorted population in each brain region. Only genes that were expressed (with CPM > 2) in at least six samples were kept. The read counts of the remaining genes were then normalized using the TMM method (*125*), and DEGs were called in the quasi-likelihood F-test mode with a cutoff of FDR < 0.05.

**snATAC-seq data preprocessing, clustering and cell type annotation**

snATAC-seq reads were mapped to the human GRCh38 genome using 10x Genomics Cell Ranger ATAC (v1.1.0) (*110*) with the default parameters. Barcode multiplets (*126*) were removed using the "clean_barcode_multiplets_1.1.py" script provided by 10x Genomics. The output fragments files were then imported into ArchR (v1.0.2) (*111*) to create Arrow files and an "ArchRProject" object for the downstream analysis. Fragment size distribution in each sample was inspected for nucleosomal periodicity, and nuclei with TSS (Transcription Start Site) enrichment score less than 4 and a number of unique fragments less than 1000 were removed. Doublet inference and removal were performed using the "addDoubletScores" and "filterDoublets" functions in ArchR. A cell-by-tile matrix containing insertion counts across genome-wide 500-bp bins was created, and dimensionality reduction was applied to the matrix using the iterative Latent Semantic Indexing (LSI) implemented in ArchR (two iterations with the top 25,000 features). The top 30 dimensions after LSI were used to perform Louvain clustering (with resolution set as 2) and UMAP visualization.

Next, a cell-by-gene matrix was computed with the gene activity score model implemented by ArchR. The model accounted for both the accessibility within the entire gene body and the activity of putative distal regulatory elements using an exponential weighting function that depends on the distance between the insertions and the TSS of the gene of interest, and gene boundaries were imposed to minimize the contribution of unrelated regulatory elements to the gene activity score. Gene activity scores of well-defined cell class marker genes were used to classify the clusters into excitatory neurons, inhibitory neurons and non-neurons. A subset of nuclei was created for each of these three classes, and a similar LSI and clustering pipeline was run to identify subclusters in each category. For visualization, the "addImputeWeights" function was used to impute gene activity scores by smoothing signals across nearby cells based on a MAGIC (*127*) diffusion matrix.

To annotate these subclusters, we integrated the gene activity score matrix in the snATAC dataset with the gene expression matrix in our snRNA dataset using the "addGeneIntegrationMatrix" function. The integration was done in a constrained way, where subclusters from the three cell classes in the snATAC dataset were only allowed to align to the subclusters with matching cell classes in the snRNA dataset. The annotation labels from the 14 major snRNA cell types were transferred to each snATAC nuclei with a prediction score to represent the assignment accuracy. The final annotations of the snATAC subclusters were determined by first filtering out nuclei with a prediction score less than 0.7 and then taking the transferred label of 70% supermajority in remaining nuclei for each cluster. Subclusters that failed to reach the supermajority were labeled as “Mixed” and removed from further consideration. This resulted in 109,198 high-quality snATAC nuclei of 11 major brain cell types.

**snATAC-seq peak calling**

To call peaks in each group of interest (the combination of major cell types, brain regions and diagnosis), we first created pseudo-bulk replicates using the “addGroupCoverages” function in ArchR with the following settings: the minimum and maximum numbers of replicates were set to 6 and 28; the minimum and maximum numbers of nuclei per replicate were set to 50 and 500; the sampling ratio to use if a particular group lacks sufficient cells to make the desired replicates was set to 0.8. Next, the “addReproduciblePeakSet” in ArchR was run with MACS2 (v2.2.7.1) (*112*) set as the peak calling method, where the iterative overlap peak merging procedure was performed to generate a single merged peak set of fixed-width (501 bp), reproducible peaks that can be called in at least two samples. A cell-by-peak count matrix was then computed with the “addPeakMatrix” function. The counts were normalized by the number of reads in TSS across cells prior to performing marker feature identification, and the cell-type-specific peaks were identified using the Wilcoxon test in the “getMarkerFeatures” function for each major cell type, adjusting for the potential bias introduced by the number of unique fragments and TSS enrichment.

**Differential chromatin accessibility between disease and control**

Differential chromatin accessibility between disease and control was accessed in two ways. First, we ran gene-based pair-wise differential tests with the gene activity scores for each major cell type using the Wilcoxon test in the “getMarkerFeatures” function, accounting for bias of the number of unique fragments and TSS enrichment. Genes with FDR less than 0.05 were called significant. In addition, peak-based differential tests were performed to identify differentially accessible regions (DARs) between disease and control for each major cell type. The cell-by-peak matrix was normalized by the number of reads in TSS across cells and DARs were identified using the Wilcoxon test in the “getMarkerFeatures” function, adjusting for the potential bias introduced by the number of unique fragments and TSS enrichment. No significant DARs were found with an FDR threshold of 0.05.

**Accessibility of transcription factor motif**

ChromVAR (*72*) was used to estimate chromatin accessibility within peaks sharing the same transcription factor (TF) motif while controlling for technical biases. First, the non-redundant transcription factor (TF) motif archetypes (v2.0-beta, https://github.com/jvierstra/motif-clustering) (*128*) were used to scan and annotate the peaks with the “addMotifAnnotations” function in ArchR. Background peaks were selected based on similarity in GC content and the number of fragments across all samples using the “addBgdPeaks” function. The chromVAR deviations and z scores per cell were then computed for each TF motif using the “addDevationsMatrix” function. To find TF motifs with differential chromVAR scores between disease and control in each major cell type, t-test and Brown-Forsythe test were performed on the deviations and z scores, respectively. Motifs with FDR < 0.05 were called significant. Motif sequence logos were drawn using the “ceqlogo” module in the MEME Suite (v5.4.1) (*129*).

**H3K27ac ChIP-seq data processing**

Raw sequencing data were pre-processed to remove adapters and low-quality sequences with the HTStream tool (https://github.com/s4hts/HTStream). Reads were mapped to the human hg38 genome build with BWA-MEM2 (*113*) and filtered to remove multi-mapping reads and low-quality alignments using Samtools (*114*). Reads mapping to ENCODE-blacklisted genomic regions were excluded using BEDTools (*115*). H3K27ac-enriched peaks were detected using MACS2, including input controls for each cell type and condition, as previously described (*116*). Promoter (2 kb upstream and 1 kb downstream from TSS) H3K27ac signal was computed using the “multiBigwigSummary” module from deepTools (v.3.3.1) (*117*). Differential peaks between C9-ALS and control for each FANS-sort population were identified using DiffBind (*118*) in edgeR (*109*) mode. Peaks with FDR < 0.05 were called significant.

#### Supplementary Figures

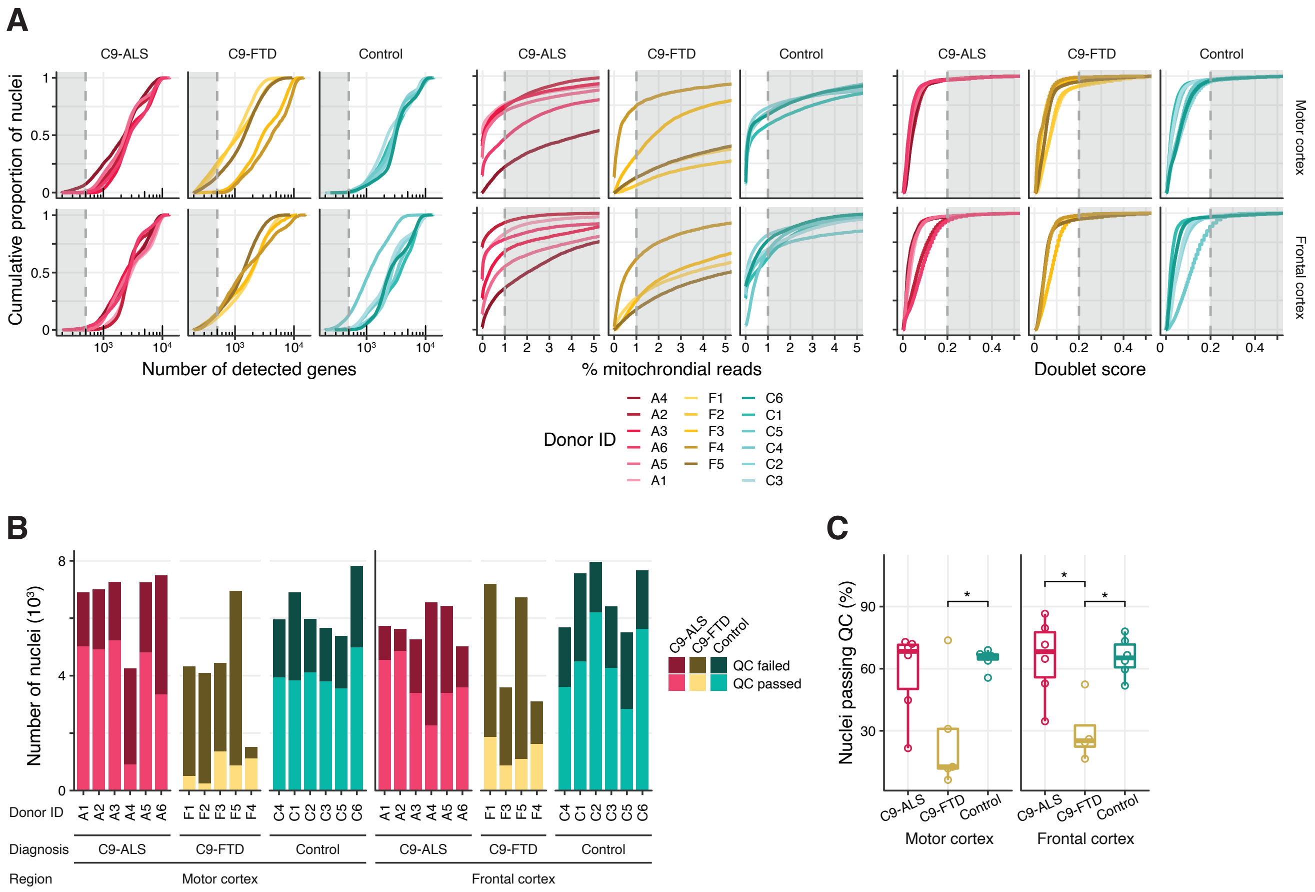


**Fig. S1. Quality control (QC) metrics for snRNA-seq data.** (A) Cumulative proportion of the number of detected genes, percentage of reads mapped to the mitochondrial genome, and doublet score (from scrublet (*100*)) for nuclei in each snRNA-seq sample, grouped by donor ID, diagnosis and brain region. Shaded areas in gray mark the nuclei that failed the QC filtering (number of detected genes > 500, % mitochondrial reads < 1, or doublet score < 0.2). Nuclei in the C9-FTD samples failed QC mainly due to high percentage of mitochondrial reads. (B) Summary of the total number of collected nuclei, and the number that passed QC, in each sample. (C) Proportion of nuclei that passed QC. Circles represent individual donors. *, t-test $p < 0.05$.

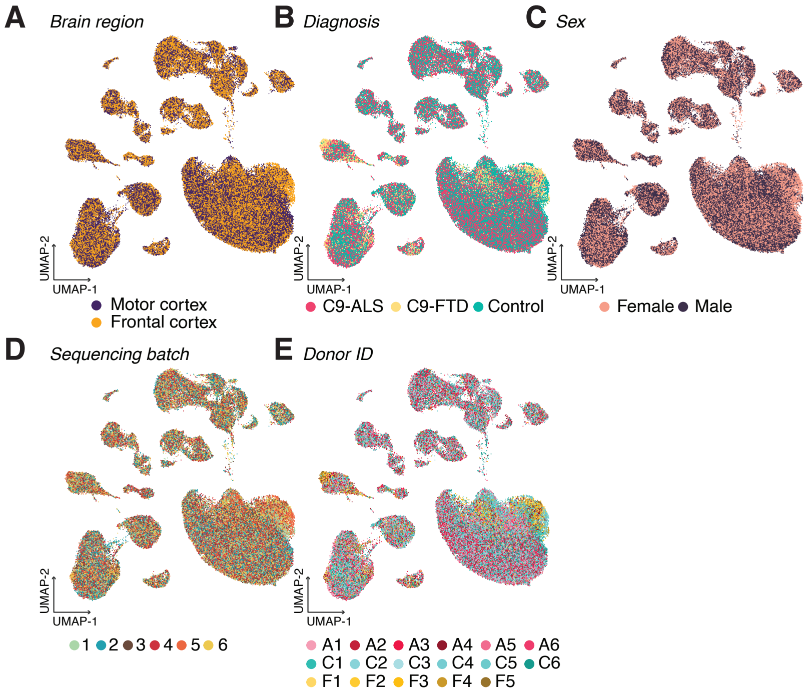


**Fig. S2. The clusters identified in snRNA-seq were not biased by known covariates.** Uniform manifold approximation and projection (UMAP, (*104*)) embedding of snRNA-seq profiles (see Fig. 1A) were colored by brain region (A), diagnosis (B), donor sex (C), sequencing batch (D), or individual donor (E).

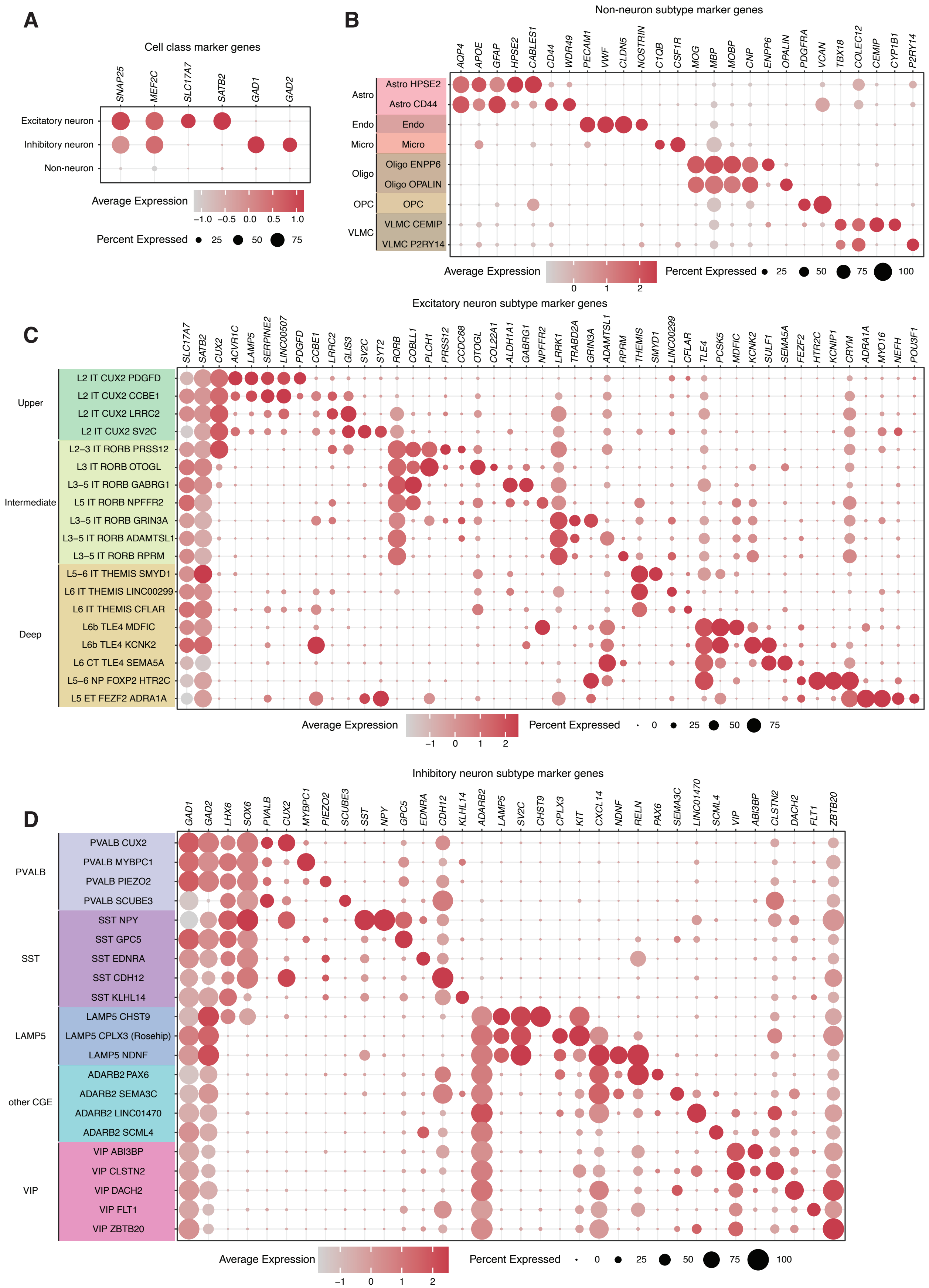
A
Cell class marker genes
SNAP25
MEF2C
SLC17A7
SATB2
GAD1
GAD2
Excitatory neuron
Inhibitory neuron
Non-neuron
Average Expression
−1.0 −0.5 0.0 0.5 1.0
Percent Expressed 25 50 75
B
Non-neuron subtype marker genes
AQP4 APOE GFAP HPSE2 CABLES1 CD44 WDR49 PECAM1 VWF CLDN5 NOSTRIN C1QB CSF1R MOG MBP MOBP CNP ENPP6 OPALIN PDGFRA VCAN TBX18 COLEC12 CEMIP CYP1B1 P2RY14
Astro
Astro HPSE2
Astro CD44
Endo
Micro
Oligo
Oligo ENPP6
Oligo OPALIN
OPC
VLMC
VLMC CEMIP
VLMC P2RY14
Average Expression
0 1 2
Percent Expressed 25 50 75 100
C
Excitatory neuron subtype marker genes
SLC17A7 SATB2 CUX2 ACVR1C LAMP5 SERPINE2 LINC00507 PDGFD CCBE1 LRRC2 GLIS3 SV2C SYT2 RORB COBLL1 PLCH1 PRSS12 CCDC68 OTOGL COL22A1 ALDH1A1 GABRG1 NPFFR2 LRRK1 TRABD2A GRIN3A ADAMTSL1 RPRM THEMIS SMYD1 LINC00299 CFLAR TLE4 PCSK5 MDFIC KCNK2 SULF1 SEMA5A FEZF2 HTR2C KCNIP1 CRYM ADRA1A MYO16 NEFH POU3F1
Upper
L2 IT CUX2 PDGFD
L2 IT CUX2 CCBE1
L2 IT CUX2 LRRC2
L2 IT CUX2 SV2C
Intermediate
L2–3 IT RORB PRSS12
L3 IT RORB OTOGL
L3–5 IT RORB GABRG1
L5 IT RORB NPFFR2
L3–5 IT RORB GRIN3A
L3–5 IT RORB ADAMTSL1
L3–5 IT RORB RPRM
Deep
L5–6 IT THEMIS SMYD1
L6 IT THEMIS LINC00299
L6 IT THEMIS CFLAR
L6b TLE4 MDFIC
L6b TLE4 KCNK2
L6 CT TLE4 SEMA5A
L5–6 NP FOXP2 HTR2C
L5 ET FEZF2 ADRA1A
Average Expression
−1 0 1 2
Percent Expressed 0 25 50 75
D
Inhibitory neuron subtype marker genes
GAD1 GAD2 LHX6 SOX6 PVALB CUX2 MYBPC1 PIEZO2 SCUBE3 SST NPY GPC5 EDNRA CDH12 KLHL14 ADARB2 LAMP5 SV2C CHST9 CPLX3 KIT CXCL14 NDNF RELN PAX6 SEMA3C LINC01470 SCML4 VIP ABI3BP CLSTN2 DACH2 FLT1 ZBTB20
PVALB
PVALB CUX2
PVALB MYBPC1
PVALB PIEZO2
PVALB SCUBE3
SST
SST NPY
SST GPC5
SST EDNRA
SST CDH12
SST KLHL14
LAMP5
LAMP5 CHST9
LAMP5 CPLX3 (Rosehip)
LAMP5 NDNF
other CGE
ADARB2 PAX6
ADARB2 SEMA3C
ADARB2 LINC01470
ADARB2 SCML4
VIP
VIP ABI3BP
VIP CLSTN2
VIP DACH2
VIP FLT1
VIP ZBTB20
Average Expression
−1 0 1 2
Percent Expressed 0 25 50 75 100

**Fig. S3. Annotations of the identified cellular populations in snRNA-seq based on the expression of known markers.** Nuclei were first categorized into excitatory and inhibitory neurons, and non-neuronal cells using cell class marker genes (A). Fine-grained cell types were then annotated using marker genes for non-neuronal cells (B), excitatory neurons (C), and inhibitory neurons (D). Major cell types (left colored bars) were defined using shared marker genes. Normalized expression is defined as the z-score of log(CPM) for each gene across all cells. Dots were colored by the average normalized expression in each cell type, and dot size represents the percentage of nuclei expressing the marker genes (with non-zero counts) in the cell type. CGE, caudal ganglionic eminence; Astro, astrocytes; Endo, endothelial cells; Micro, microglia; Oligo, oligodendrocytes; OPC, oligodendrocyte precursor cells; VLMC, vascular leptomeningeal cell; IT, intratelencephalic; CT, corticothalamic; NP, near-projecting; ET, extratelencephalic.

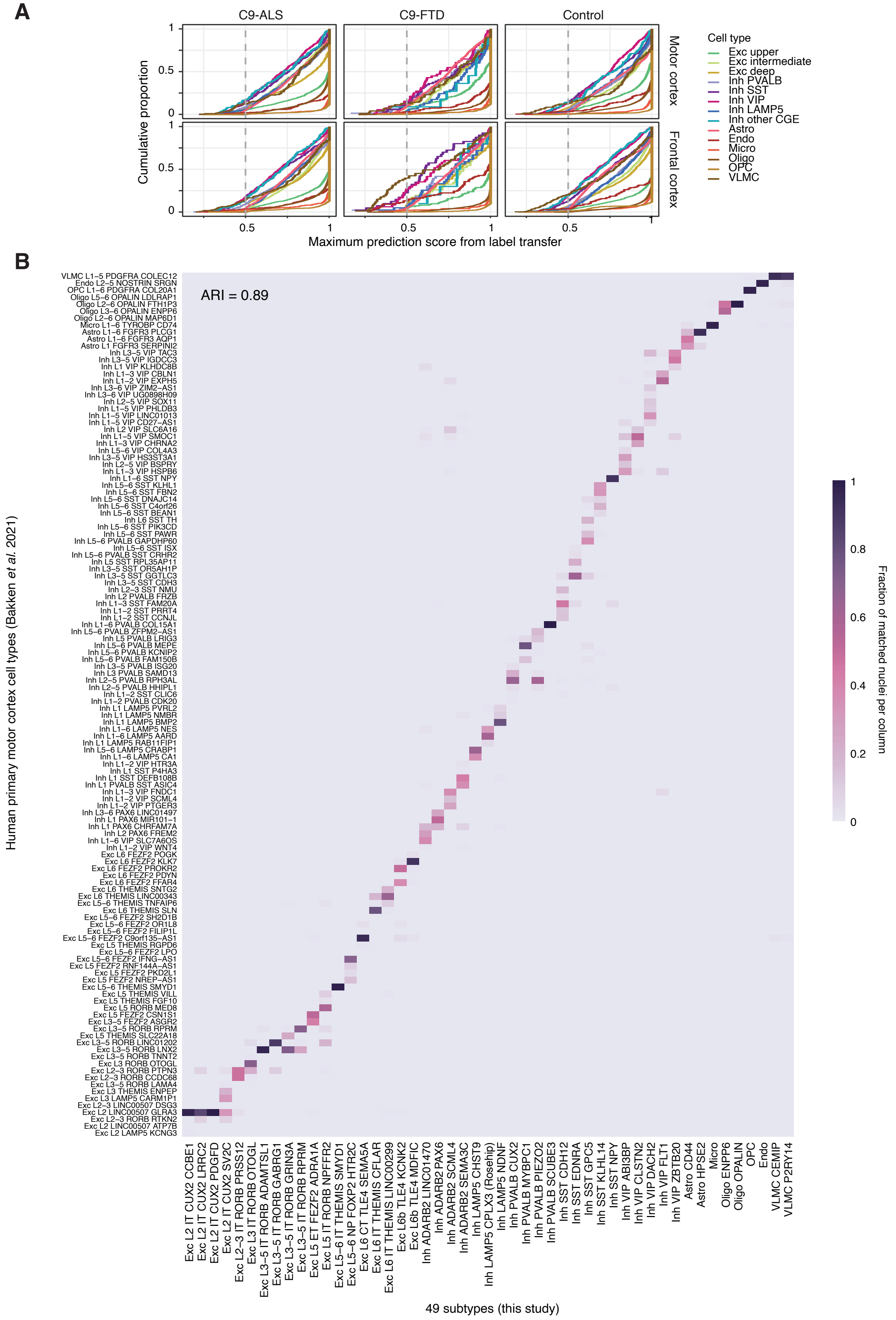
A
C9-ALS
C9-FTD
Control
Motor cortex
Frontal cortex
Cumulative proportion
Maximum prediction score from label transfer
Cell type
Exc upper
Exc intermediate
Exc deep
Inh PVALB
Inh SST
Inh VIP
Inh LAMP5
Inh other CGE
Astro
Endo
Micro
Oligo
OPC
VLMC
B
ARI = 0.89
Human primary motor cortex cell types (Bakken et al. 2021)
49 subtypes (this study)
Fraction of matched nuclei per column
VLMC L1–5 PDGFRA COLEC12
Endo L2–5 NOSTRIN SRGN
OPC L1–6 PDGFRA COL20A1
Oligo L5–6 OPALIN LDLRAP1
Oligo L2–6 OPALIN FTH1P3
Oligo L3–6 OPALIN ENPP6
Oligo L2–6 OPALIN MAP6D1
Micro L1–6 TYROBP CD74
Astro L1–6 FGFR3 PLCG1
Astro L1–6 FGFR3 AQP1
Astro L1 FGFR3 SERPINI2
Inh L3–5 VIP TAC3
Inh L3–5 VIP IGDCC3
Inh L1 VIP KLHDC8B
Inh L1–3 VIP CBLN1
Inh L1–2 VIP EXPH5
Inh L3–6 VIP ZIM2–AS1
Inh L3–6 VIP UG0898H09
Inh L2–5 VIP SOX11
Inh L1–5 VIP PHLDB3
Inh L1–5 VIP LINC01013
Inh L1–5 VIP CD27–AS1
Inh L2 VIP SLC6A16
Inh L1–5 VIP SMOC1
Inh L1–3 VIP CHRNA2
Inh L5–6 VIP COL4A3
Inh L3–5 VIP HS3ST3A1
Inh L2–5 VIP BSPRY
Inh L1–3 VIP HSPB6
Inh L1–6 SST NPY
Inh L5–6 SST KLHL1
Inh L5–6 SST FBN2
Inh L5–6 SST DNAJC14
Inh L5–6 SST C4orf26
Inh L5–6 SST BEAN1
Inh L6 SST TH
Inh L5–6 SST PIK3CD
Inh L5–6 SST PAWR
Inh L5–6 PVALB GAPDHP60
Inh L5–6 SST ISX
Inh L5–6 PVALB SST CRHR2
Inh L5 SST RPL35AP11
Inh L3–5 SST OR5AH1P
Inh L3–5 SST GGTLC3
Inh L3–5 SST CDH3
Inh L2–3 SST NMU
Inh L2 PVALB FRZB
Inh L1–3 SST FAM20A
Inh L1–2 SST PRRT4
Inh L1–2 SST CCNJL
Inh L1–6 PVALB COL15A1
Inh L5–6 PVALB ZFPM2–AS1
Inh L5 PVALB LRIG3
Inh L5–6 PVALB MEPE
Inh L5–6 PVALB KCNIP2
Inh L5–6 PVALB FAM150B
Inh L3–5 PVALB ISG20
Inh L3 PVALB SAMD13
Inh L2–5 PVALB RPH3AL
Inh L2–5 PVALB HHIPL1
Inh L1–2 SST CLIC6
Inh L1–2 PVALB CDK20
Inh L1 LAMP5 PVRL2
Inh L1 LAMP5 NMBR
Inh L1 LAMP5 BMP2
Inh L1–6 LAMP5 NES
Inh L1–6 LAMP5 AARD
Inh L1 LAMP5 RAB11FIP1
Inh L5–6 LAMP5 CRABP1
Inh L1–6 LAMP5 CA1
Inh L1–2 VIP HTR3A
Inh L1 SST P4HA3
Inh L1 SST DEFB108B
Inh L1 PVALB SST ASIC4
Inh L1–3 VIP FNDC1
Inh L1–2 VIP SCML4
Inh L1–2 VIP PTGER3
Inh L3–6 PAX6 LINC01497
Inh L1 PAX6 MIR101–1
Inh L1 PAX6 CHRFAM7A
Inh L2 PAX6 FREM2
Inh L1–6 VIP SLC7A6OS
Inh L1–2 VIP WNT4
Exc L6 FEZF2 POGK
Exc L6 FEZF2 KLK7
Exc L6 FEZF2 PROKR2
Exc L6 FEZF2 PDYN
Exc L6 FEZF2 FFAR4
Exc L6 THEMIS SNTG2
Exc L6 THEMIS LINC00343
Exc L5–6 THEMIS TNFAIP6
Exc L6 THEMIS SLN
Exc L5–6 FEZF2 SH2D1B
Exc L5–6 FEZF2 OR1L8
Exc L5–6 FEZF2 FILIP1L
Exc L5–6 FEZF2 C9orf135–AS1
Exc L5 THEMIS RGPD6
Exc L5–6 FEZF2 LPO
Exc L5–6 FEZF2 IFNG–AS1
Exc L5 FEZF2 RNF144A–AS1
Exc L5 FEZF2 PKD2L1
Exc L5 FEZF2 NREP–AS1
Exc L5–6 THEMIS SMYD1
Exc L5 THEMIS VILL
Exc L5 THEMIS FGF10
Exc L5 RORB MED8
Exc L5 FEZF2 CSN1S1
Exc L3–5 FEZF2 ASGR2
Exc L3–5 RORB RPRM
Exc L5 THEMIS SLC22A18
Exc L3–5 RORB LINC01202
Exc L3–5 RORB LNX2
Exc L3–5 RORB TNNT2
Exc L3 RORB OTOGL
Exc L2–3 RORB PTPN3
Exc L2–3 RORB CCDC68
Exc L3–5 RORB LAMA4
Exc L3 THEMIS ENPEP
Exc L3 LAMP5 CARM1P1
Exc L2–3 LINC00507 DSG3
Exc L2 LINC00507 GLRA3
Exc L2–3 RORB RTKN2
Exc L2 LINC00507 ATP7B
Exc L2 LAMP5 KCNG3
Exc L2 IT CUX2 CCBE1
Exc L2 IT CUX2 LRRC2
Exc L2 IT CUX2 PDGFD
Exc L2 IT CUX2 SV2C
Exc L2–3 IT RORB PRSS12
Exc L3 IT RORB OTOGL
Exc L3–5 IT RORB ADAMTSL1
Exc L3–5 IT RORB GABRG1
Exc L3–5 IT RORB GRIN3A
Exc L3–5 IT RORB RPRM
Exc L5 ET FEZF2 ADRA1A
Exc L5 IT RORB NPFFR2
Exc L5–6 IT THEMIS SMYD1
Exc L5–6 NP FOXP2 HTR2C
Exc L6 CT TLE4 SEMA5A
Exc L6 IT THEMIS CFLAR
Exc L6 IT THEMIS LINC00299
Exc L6b TLE4 KCNK2
Exc L6b TLE4 MDFIC
Inh ADARB2 LINC01470
Inh ADARB2 PAX6
Inh ADARB2 SCML4
Inh ADARB2 SEMA3C
Inh LAMP5 CHST9
Inh LAMP5 CPLX3 (Rosehip)
Inh LAMP5 NDNF
Inh PVALB CUX2
Inh PVALB MYBPC1
Inh PVALB PIEZO2
Inh PVALB SCUBE3
Inh SST CDH12
Inh SST EDNRA
Inh SST GPC5
Inh SST KLHL14
Inh SST NPY
Inh VIP ABI3BP
Inh VIP CLSTN2
Inh VIP DACH2
Inh VIP FLT1
Inh VIP ZBTB20
Astro CD44
Astro HPSE2
Micro
Oligo ENPP6
Oligo OPALIN
OPC
Endo
VLMC CEMIP
VLMC P2RY14

**Fig. S4. Comparison of the cell types identified in our snRNA-seq dataset with a previously published dataset.** (A) Cumulative proportion of the maximum prediction score across nuclei from the label transfer analysis using the published human primary motor cortex cell types (*24*) as the reference dataset (see Methods). Curves are grouped and colored by diagnosis, brain region, and the major cell types identified in our study. (B) The annotations in Fig.1 and Fig. S3 were consistent with cell types in the study of the human motor cortex (*24*). ARI, adjusted Rand index.

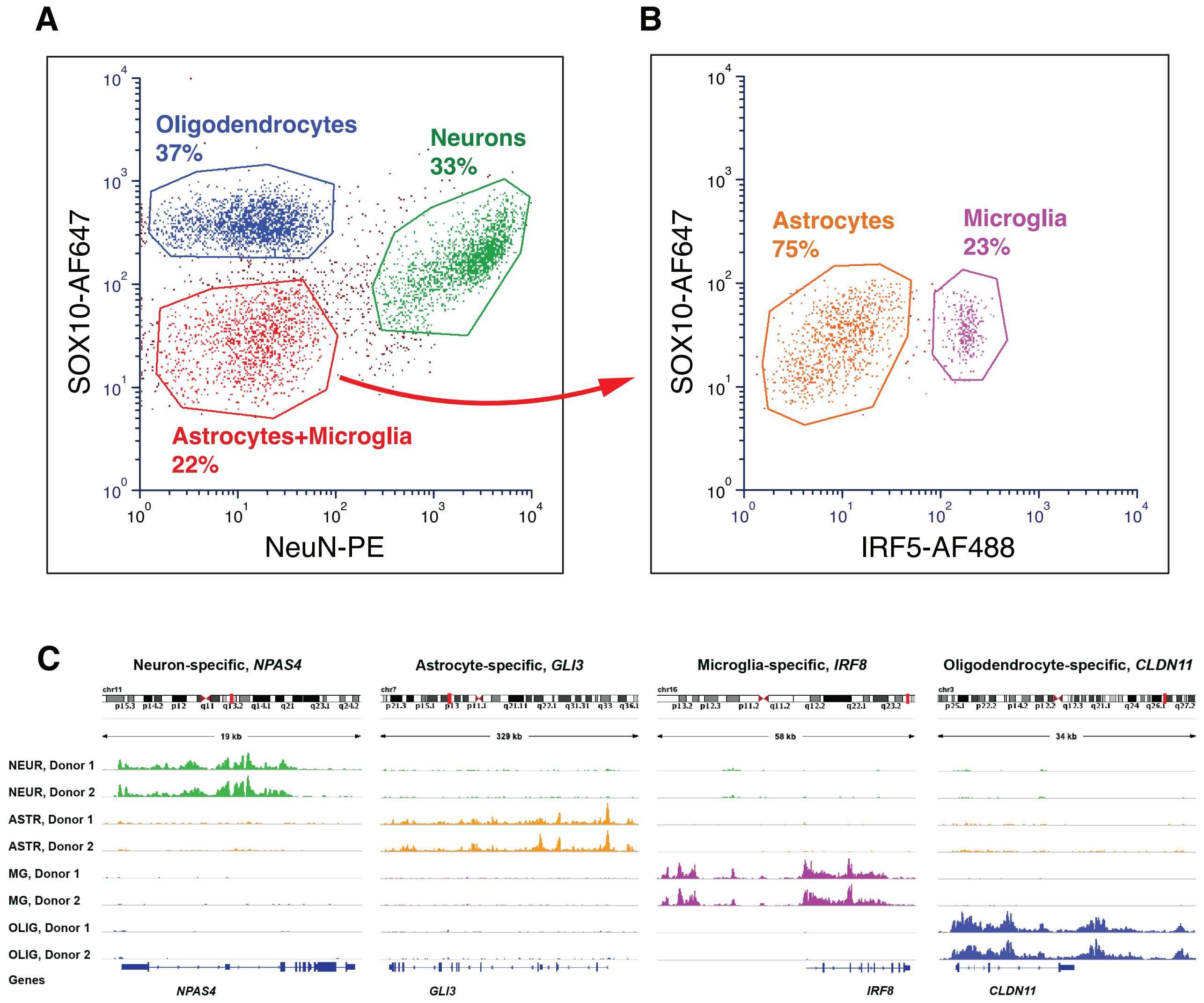


**Fig. S5. Fluorescence-activated nuclei sorting (FANS) isolation of nuclei of major brain cell types.** (A,B) Sequential FANS gating procedure to isolate nuclei from cell populations. (A) Three cell populations were isolated for bulk RNA-seq (Figs. 2E-F and 6F): neurons (NeuN+), oligodendrocyte lineage cells (NeuN–/Sox10+, consisting mainly of mature oligodendrocytes and a smaller population of OPCs), and other glia (NeuN–/Sox10–, mostly consisting of microglia and astrocytes). Anti-NeuN and anti-SOX10 antibodies were used to separate these three populations. (B) For H3K27ac ChIP-seq (Fig. 5), the NeuN–/Sox10– population was further split to isolate microglia (IRF5+), and astrocytes (IRF5–). (C) Validation of FANS-separated neurons, oligodendrocyte lineage cells, microglia and astrocytes using H3K27ac ChIP-seq. Signals from two donors are shown. In each population, H3K27ac signal enrichment was detected for known cell-type-specific genes. NEUR, neurons; ASTR, astrocytes; OLIG, oligodendrocyte lineage; MG, microglia.

A

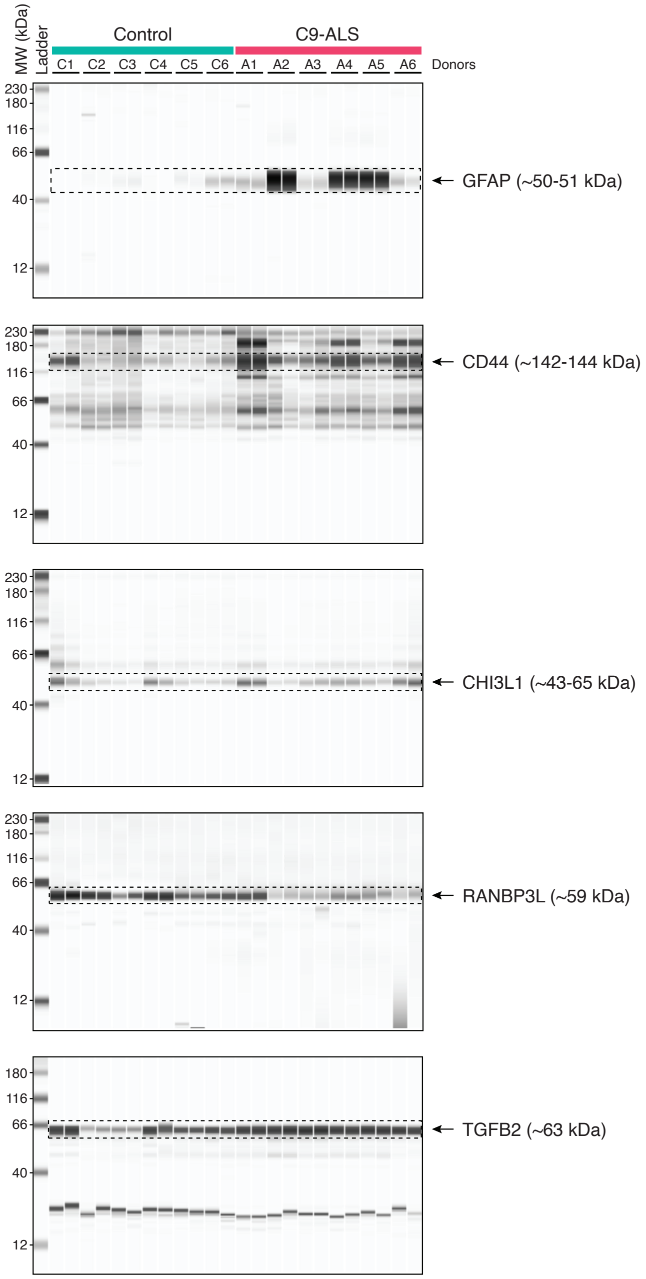


**Fig. S6. Quantification of proteins encoded by genes dysregulated in C9-ALS astrocytes using automated Western blot analysis.** (A) Protein levels were determined in selected DE genes identified in astrocytes in our snRNA-seq analysis. Bulk motor cortex tissues (~80-100mg) from six C9-ALS and six control donors were used. Each sample was assessed in duplicate. Automated capillary Western blot analysis was performed using the ProteinSimple Jess-Wes System (Methods). See Fig. 3C for the quantifications of protein levels. A two-sided Welch's t-test was used to compare the immunoreactive signals between C9-ALS and control, considering the averaged signals between two replicates for each donor as an observation. GFAP, CD44, and TGFB2 proteins were upregulated ($p = 0.033, 0.012, 0.033$ respectively) in C9-ALS motor cortex, whereas RANBP3L protein trended in the expected direction ($p = 0.063$, respectively) and CHI3L1 was not significant ($p = 0.303$).

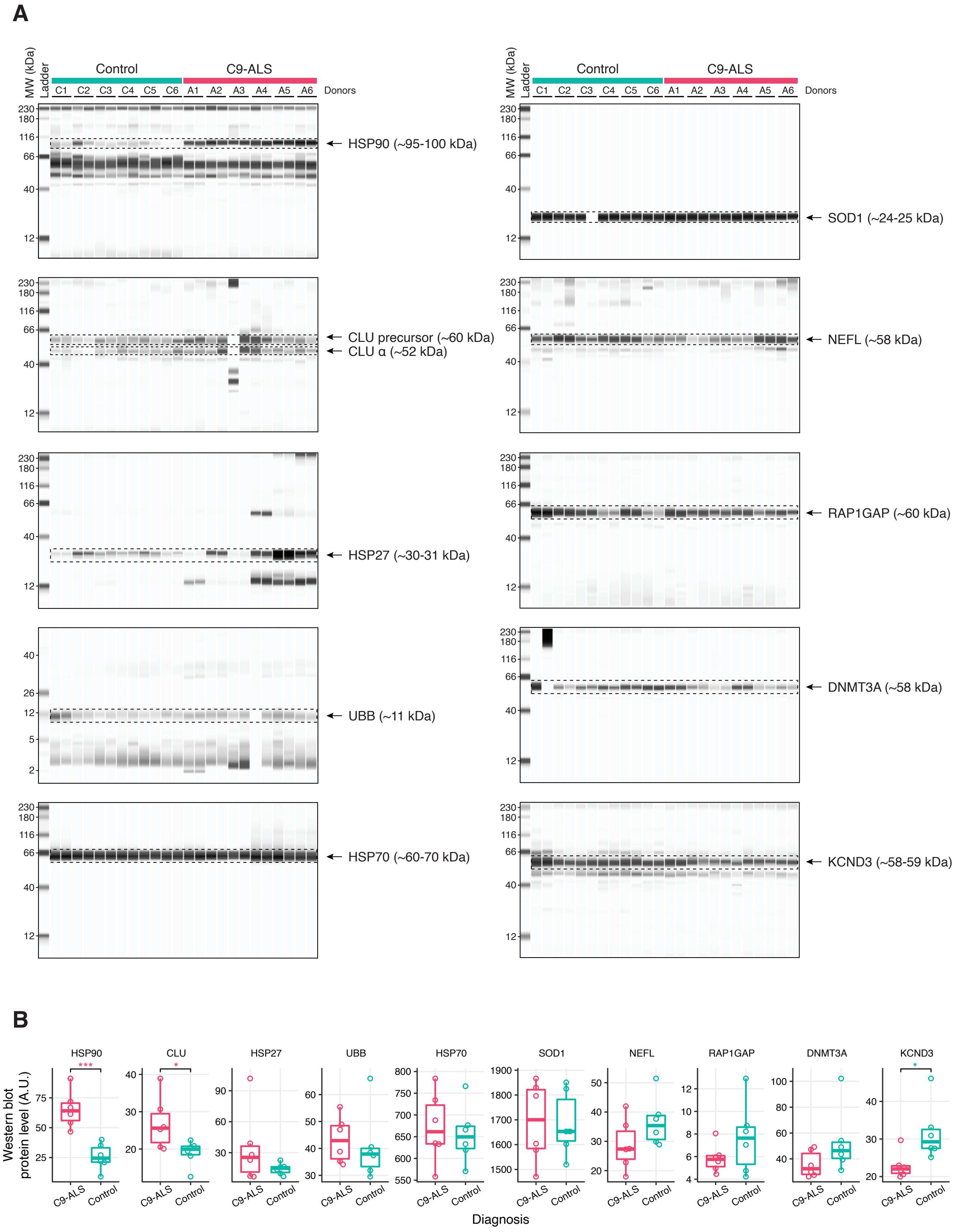
A
MW (kDa)
Ladder
Control
C9-ALS
C1 C2 C3 C4 C5 C6 A1 A2 A3 A4 A5 A6
Donors
HSP90 (~95-100 kDa)
SOD1 (~24-25 kDa)
CLU precursor (~60 kDa)
CLU α (~52 kDa)
NEFL (~58 kDa)
HSP27 (~30-31 kDa)
RAP1GAP (~60 kDa)
UBB (~11 kDa)
DNMT3A (~58 kDa)
HSP70 (~60-70 kDa)
KCND3 (~58-59 kDa)
B
HSP90
CLU
HSP27
UBB
HSP70
SOD1
NEFL
RAP1GAP
DNMT3A
KCND3
Western blot protein level (A.U.)
C9-ALS
Control
Diagnosis

**Fig. S7. Quantification of proteins encoded by genes dysregulated in C9-ALS excitatory neurons using automated Western blot analysis.** (A) Same as Fig. S6A but for selected DE genes identified in excitatory neurons with our snRNA-seq dataset. (B) Quantification of protein levels in (A) with automated Western blot analysis. Circles represent the average signal across duplicates for each donor. *, two-sided Welch's t-test $p < 0.05$; ***, $p < 0.001$. HSP90 and CLU proteins were upregulated ($p = 6.35e-4$ and 0.041, respectively), KCND3 was downregulated ($p = 0.036$), and HSP27, RAP1GAP and DNMT3A proteins were not significant ($p = 0.226$, 0.238 and 0.163, respectively). None of the tested genes had a change in protein abundance in the opposite direction from the mRNA expression change.

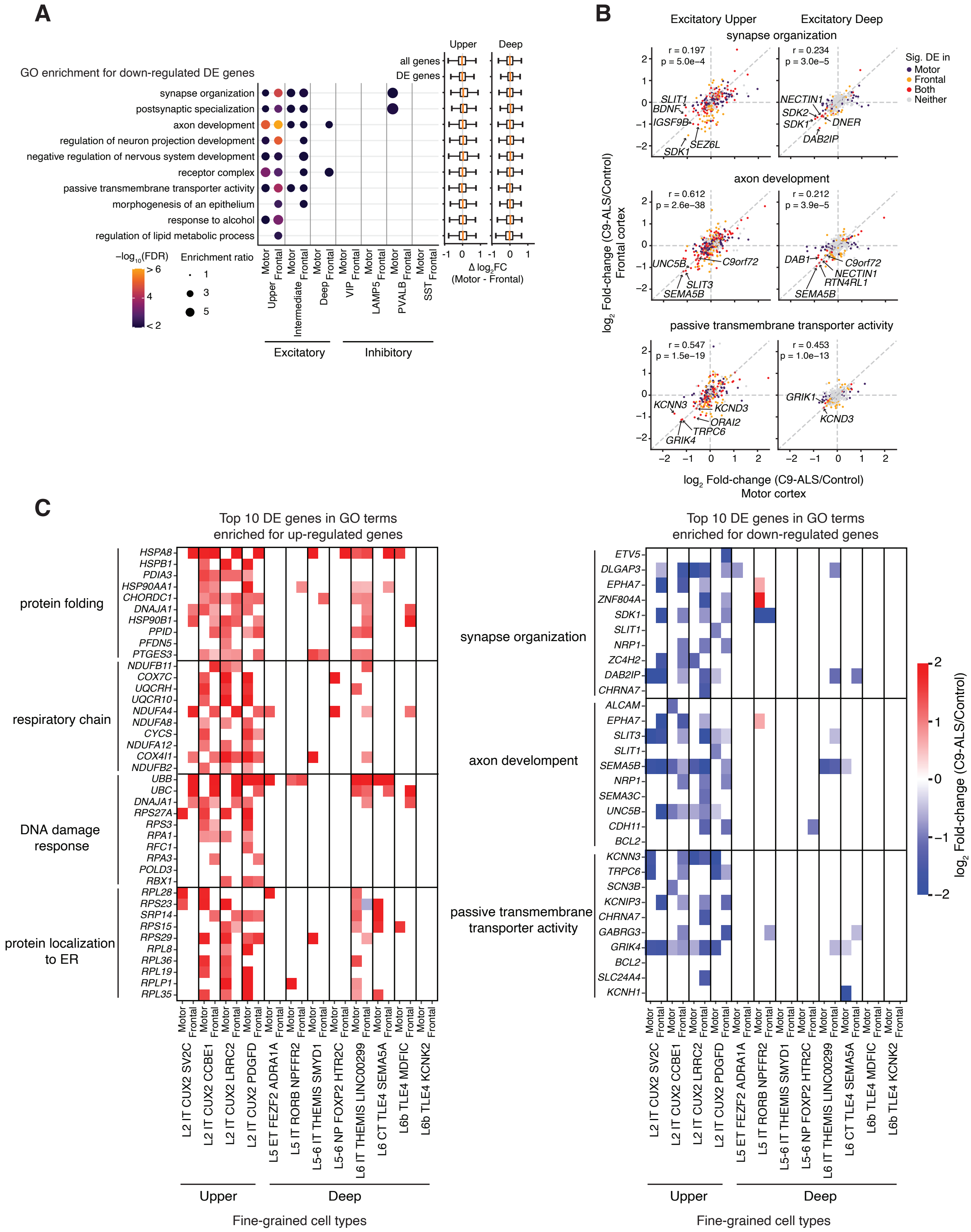
A
GO enrichment for down-regulated DE genes
all genes
DE genes
synapse organization
postsynaptic specialization
axon development
regulation of neuron projection development
negative regulation of nervous system development
receptor complex
passive transmembrane transporter activity
morphogenesis of an epithelium
response to alcohol
regulation of lipid metabolic process
Upper
Deep
−log10(FDR)
> 6
4
< 2
Enrichment ratio
1
3
5
Motor
Frontal
Upper
Intermediate
Deep
VIP
LAMP5
PVALB
SST
Excitatory
Inhibitory
Δ log2FC
(Motor - Frontal)
B
Excitatory Upper
Excitatory Deep
synapse organization
r = 0.197
p = 5.0e−4
r = 0.234
p = 3.0e−5
SLIT1
BDNF
IGSF9B
SEZ6L
SDK1
NECTIN1
SDK2
SDK1
DNER
DAB2IP
Sig. DE in
Motor
Frontal
Both
Neither
axon development
r = 0.612
p = 2.6e−38
r = 0.212
p = 3.9e−5
UNC5B
C9orf72
SLIT3
SEMA5B
DAB1
C9orf72
NECTIN1
RTN4RL1
SEMA5B
passive transmembrane transporter activity
r = 0.547
p = 1.5e−19
r = 0.453
p = 1.0e−13
KCNN3
KCND3
ORAI2
TRPC6
GRIK4
GRIK1
KCND3
log2 Fold-change (C9-ALS/Control)
Frontal cortex
log2 Fold-change (C9-ALS/Control)
Motor cortex
C
Top 10 DE genes in GO terms
enriched for up-regulated genes
protein folding
HSPA8
HSPB1
PDIA3
HSP90AA1
CHORDC1
DNAJA1
HSP90B1
PPID
PFDN5
PTGES3
respiratory chain
NDUFB11
COX7C
UQCRH
UQCR10
NDUFA4
NDUFA8
CYCS
NDUFA12
COX4I1
NDUFB2
DNA damage
response
UBB
UBC
DNAJA1
RPS27A
RPS3
RPA1
RFC1
RPA3
POLD3
RBX1
protein localization
to ER
RPL28
RPS23
SRP14
RPS15
RPS29
RPL8
RPL36
RPL19
RPLP1
RPL35
Top 10 DE genes in GO terms
enriched for down-regulated genes
synapse organization
ETV5
DLGAP3
EPHA7
ZNF804A
SDK1
SLIT1
NRP1
ZC4H2
DAB2IP
CHRNA7
axon develompent
ALCAM
EPHA7
SLIT3
SLIT1
SEMA5B
NRP1
SEMA3C
UNC5B
CDH11
BCL2
passive transmembrane
transporter activity
KCNN3
TRPC6
SCN3B
KCNIP3
CHRNA7
GABRG3
GRIK4
BCL2
SLC24A4
KCNH1
log2 Fold-change (C9-ALS/Control)
2
1
0
−1
−2
Motor
Frontal
L2 IT CUX2 SV2C
L2 IT CUX2 CCBE1
L2 IT CUX2 LRRC2
L2 IT CUX2 PDGFD
L5 ET FEZF2 ADRA1A
L5 IT RORB NPFFR2
L5-6 IT THEMIS SMYD1
L5-6 NP FOXP2 HTR2C
L6 IT THEMIS LINC00299
L6 CT TLE4 SEMA5A
L6b TLE4 MDFIC
L6b TLE4 KCNK2
Upper
Deep
Fine-grained cell types

**Fig. S8. Functional enrichment analysis of differentially expressed (DE) genes in upper- and deep-layer excitatory neurons.** (A) *Left panel*: Top gene ontology (GO) terms enriched for genes downregulated in upper and/or deep layer excitatory neurons. Enrichment of the same terms for downregulated DE genes in other neuronal cell types are shown for comparison. Enriched GO categories (FDR<0.01) were selected by affinity propagation. *Right panel*: Difference in the fold-change ($\Delta \log_2 FC$) between motor and frontal cortex for all expressed genes, all DE genes, and DE genes in each GO category. T-test was used to test whether the $\Delta \log_2 FC$ in each group of genes were significantly different from the $\Delta \log_2 FC$ of the "all genes" control set. (B) Comparison of effects in motor vs. frontal cortices for GO categories exemplifying processes/structures that are specific for neurons (*synapse organization*, *axon development*), or important for neuronal function (*passive transmembrane transport activity*). r, Pearson correlation coefficient. (C) mRNA expression fold-change (C9-ALS vs. control) of top 10 DE genes for GO terms enriched for upregulated DE genes (left panel) and downregulated genes (right panel) in fine-grained upper- and deep-layer excitatory neurons. See Table S5 for the full list of GO enrichment results.

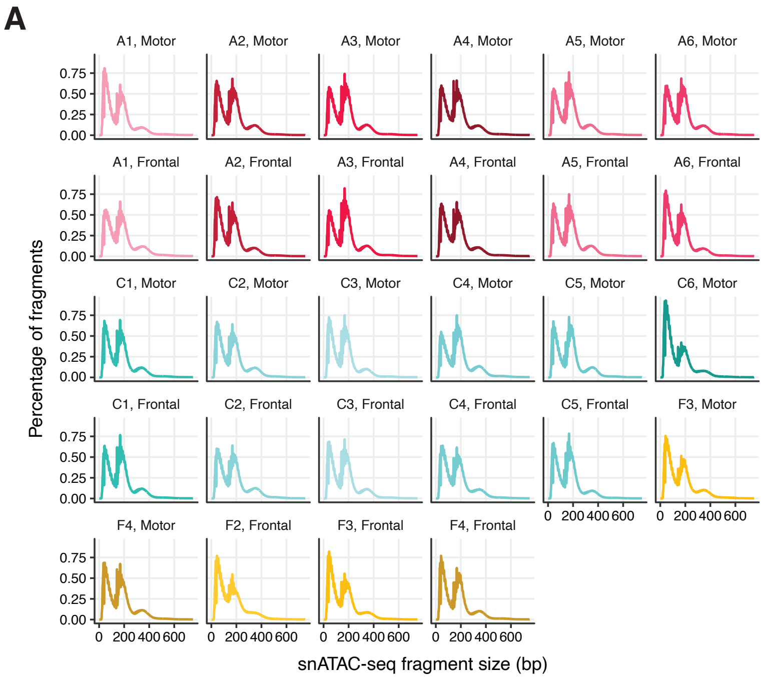


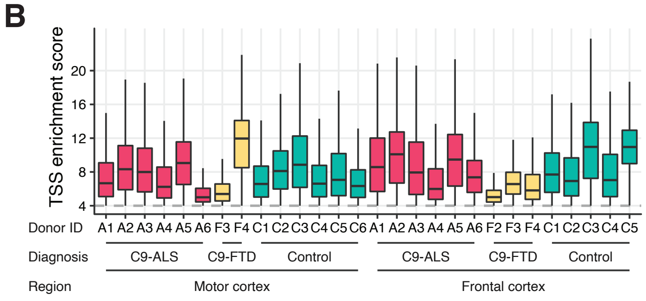


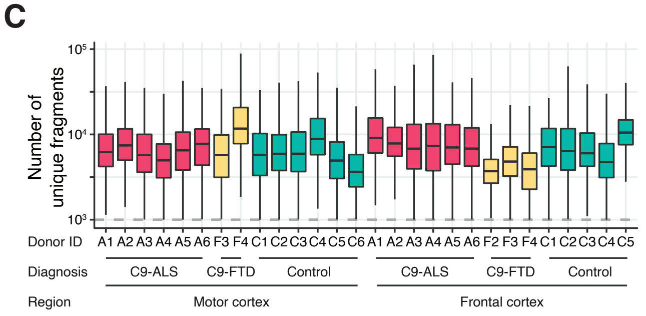


**Fig. S9. QC metrics used in the snATAC-seq data pre-processing.** (A) Distribution of snATAC-seq fragment sizes in each sample that passed QC, showing periodicity related to nucleosome spacing. (B-C) Box-and-whisker plots to show the distribution of transcription start site (TSS) enrichment score (B) and number of unique fragments (C) across nuclei grouped by samples.

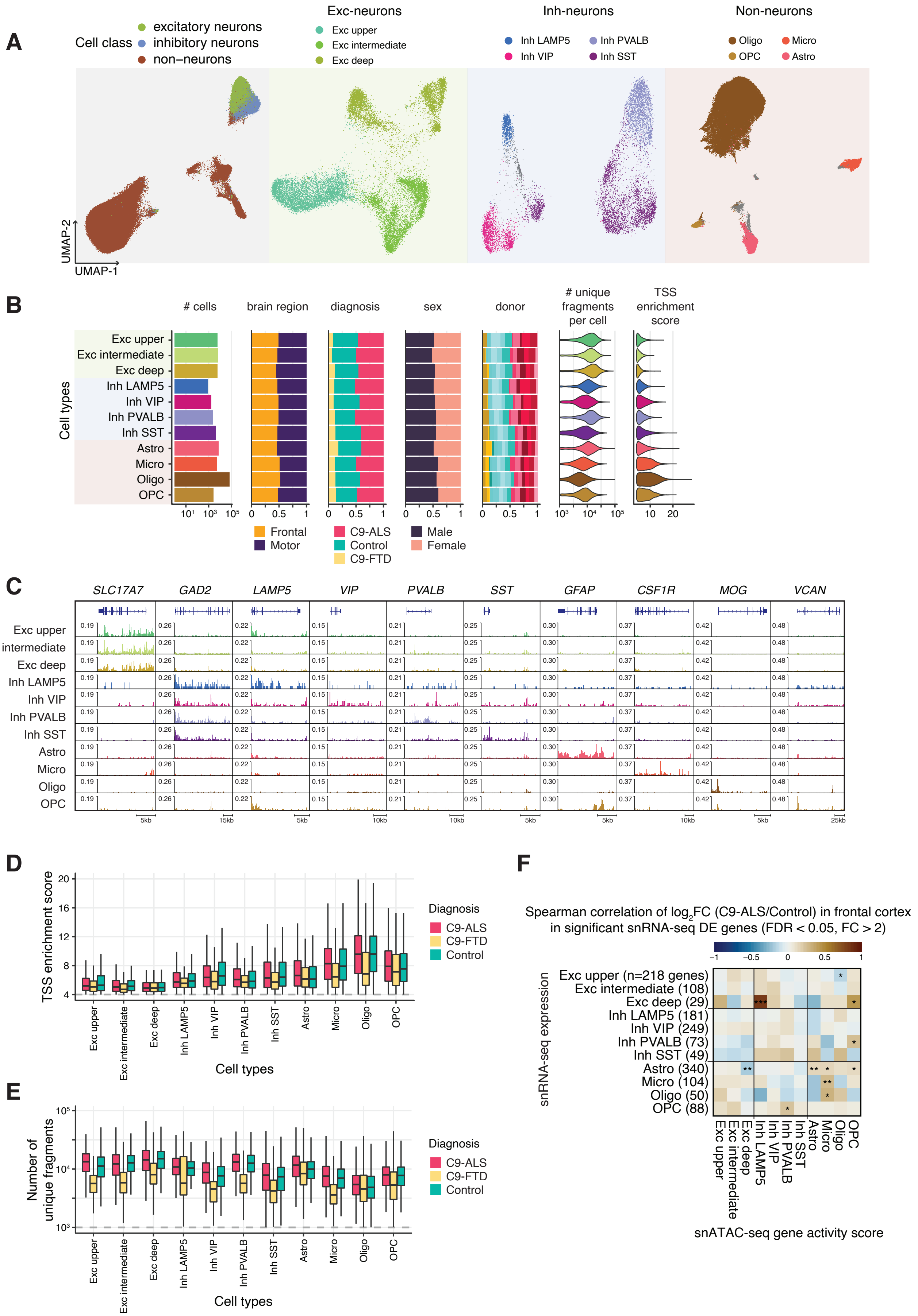
A
Cell class
excitatory neurons
inhibitory neurons
non–neurons
Exc-neurons
Exc upper
Exc intermediate
Exc deep
Inh-neurons
Inh LAMP5
Inh VIP
Inh PVALB
Inh SST
Non-neurons
Oligo
OPC
Micro
Astro
UMAP-2
UMAP-1
B
### cells
brain region
diagnosis
sex
donor
### unique fragments per cell
TSS enrichment score
Cell types
Frontal
Motor
C9-ALS
Control
C9-FTD
Male
Female
C
SLC17A7
GAD2
LAMP5
VIP
PVALB
SST
GFAP
CSF1R
MOG
VCAN
D
TSS enrichment score
Cell types
Diagnosis
C9-ALS
C9-FTD
Control
E
Number of unique fragments
Cell types
F
Spearman correlation of log2FC (C9-ALS/Control) in frontal cortex in significant snRNA-seq DE genes (FDR < 0.05, FC > 2)
Exc upper (n=218 genes)
Exc intermediate (108)
Exc deep (29)
Inh LAMP5 (181)
Inh VIP (249)
Inh PVALB (73)
Inh SST (49)
Astro (340)
Micro (104)
Oligo (50)
OPC (88)
snRNA-seq expression
snATAC-seq gene activity score

**Fig. S10. The epigenetic landscape of C9-ALS brain cells determined by snATAC-seq**. (A) Clustering n=109,198 high-quality snATAC-seq profiles (TSS enrichment ≥ 4, unique fragments ≥ 1,000 per cell) identified 11 major brain cell types. The clusters were annotated by transferring labels from the snRNA-seq data (see Methods). (B) Major cell types from snATAC-seq were distributed across brain regions, diagnosis groups, sex, and donors. TSS, transcription start site. (C) snATAC-seq signal at cell-type-specific marker genes. Track height represents pseudo-bulk counts normalized by reads in TSS. (D-E) Box-and-whisker plots to show the distribution of TSS enrichment score (D) and number of unique fragments (E) across nuclei grouped by major cell types. (F) Spearman correlation of the C9-ALS vs. control fold-change (FC) for snRNA expression vs. snATAC gene activity score in frontal cortex; the corresponding analysis for motor cortex is shown in Fig. 5E. The analysis was performed for strongly DE genes (FC > 2) in each major cell type in frontal cortex. *, $p < 0.05$; **, $p < 0.01$; ***, $p < 0.001$.

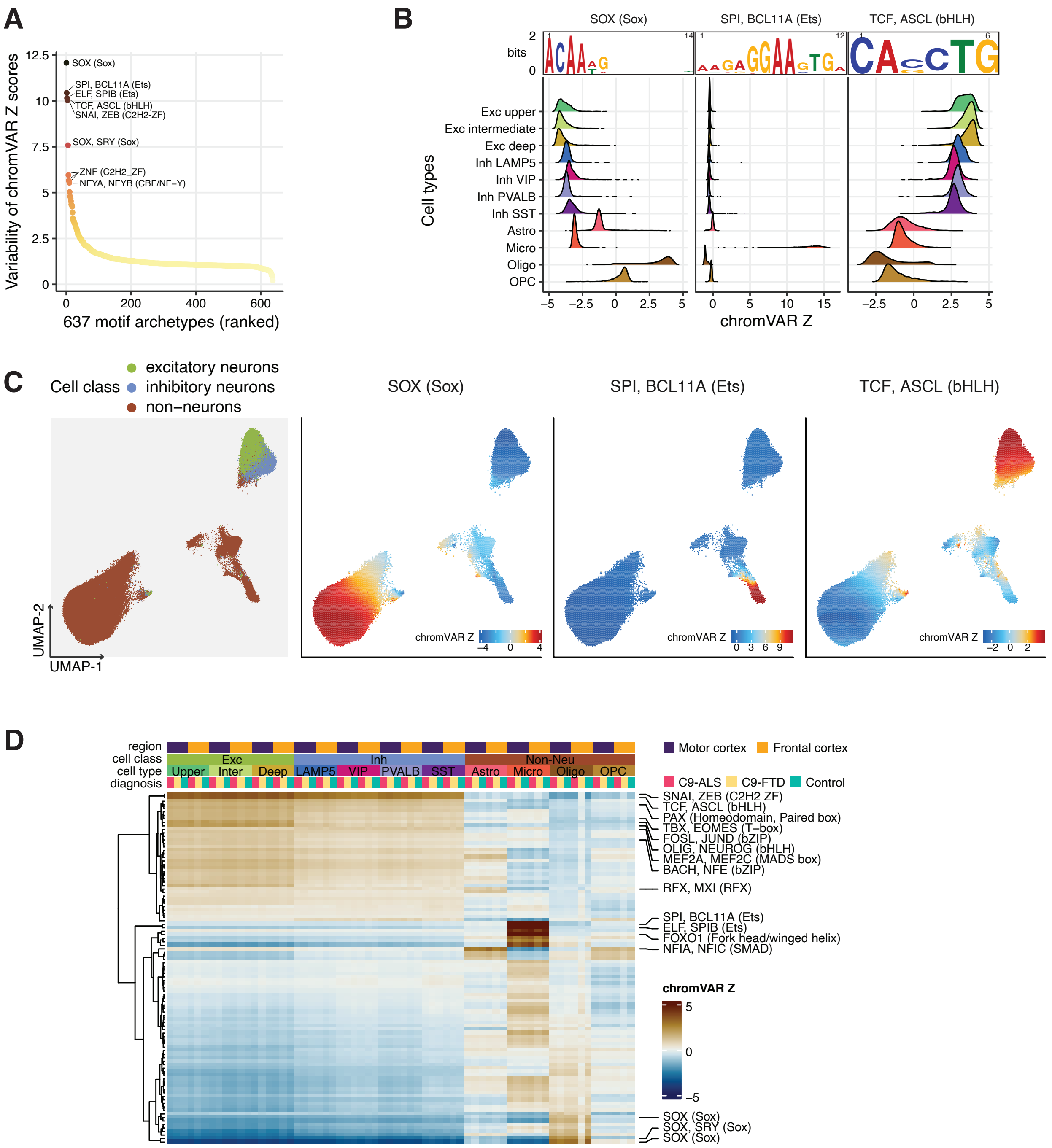


**Fig. S11. ChromVAR scores in non-redundant transcription factor (TF) motif archetypes.** (A) Variability (standard deviation) of the ChromVAR Z scores across all nuclei for TF motif archetypes (see Methods; (*72*)). (B) Distribution of ChromVAR Z score across nuclei from each major cell type for three representative TF motif archetypes with high variability. Sequence logos of the motif archetypes are shown on top. (C) ChromVAR Z scores of motif archetypes for each cell are displayed on the UMAP embedding. (D) Heatmap of average ChromVAR Z scores across groups of nuclei (columns) by major cell types, brain region and diagnosis, for the top 100 motif archetypes (rows) with highest variability across all nuclei. Top 3 cell-type-specific motif archetypes for each cell type are labeled.

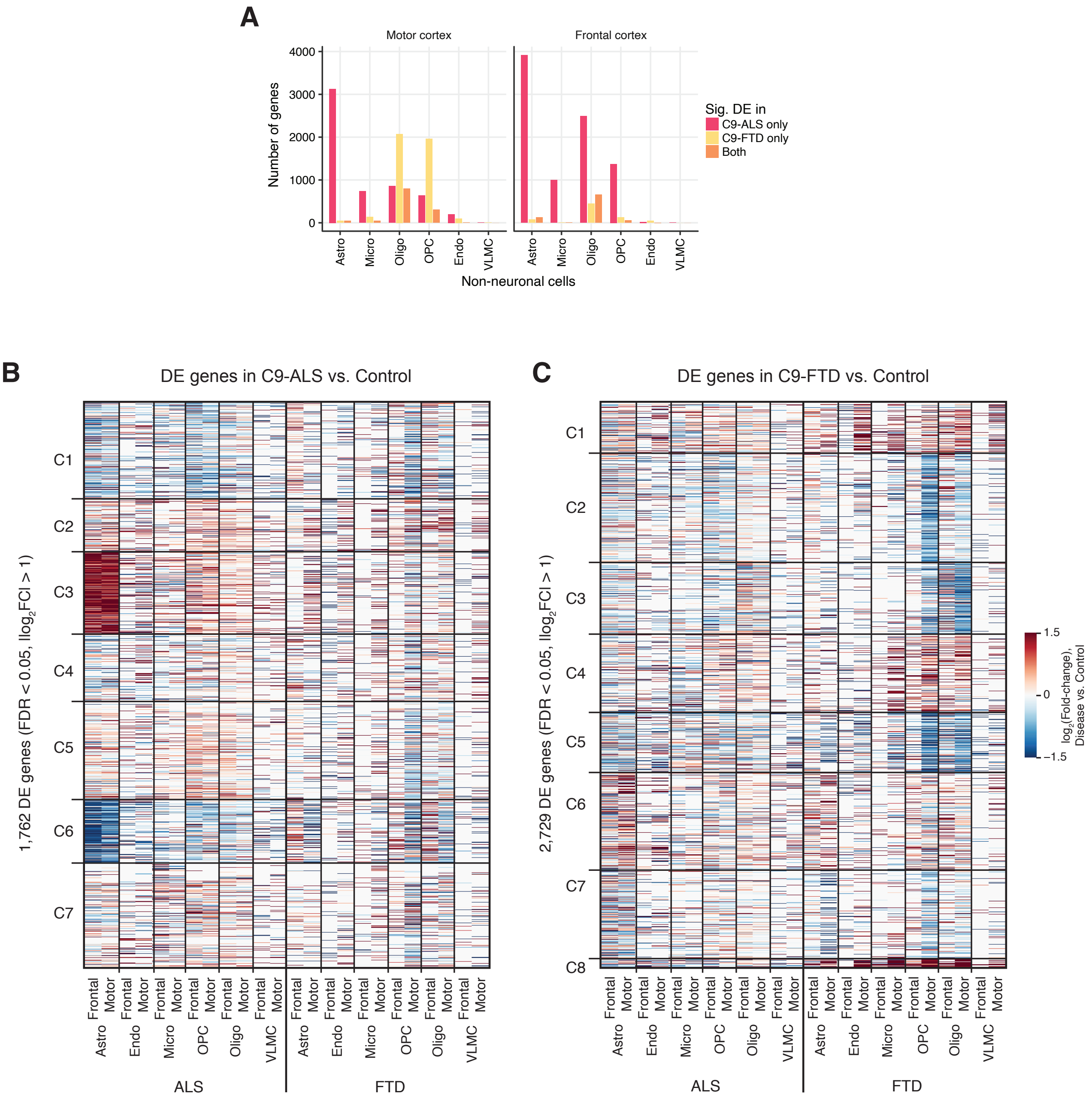


**Fig. S12. Comparison of the disease effect on gene expression in non-neuronal cells in C9-ALS and C9-FTD.** (A) Number of significant DE genes identified in the two diseases. For this analysis, control samples were split into two independent groups (from different donors) for comparison with C9-ALS and with C9-FTD to avoid double dipping (see Methods). (B-C) Heatmap to compare the gene expression fold-changes of C9-ALS vs. control and C9-FTD vs. control. Strongly DE genes (fold-change > 2) identified from the full dataset (same genes shown in Fig. 2D for C9-ALS and Fig. 6H for C9-FTD) were used, and the fold-changes from the split control analysis were shown to avoid double dipping.
